# Structure, disorder, and dynamics in task-trained recurrent neural circuits

**DOI:** 10.64898/2026.03.02.708943

**Authors:** David G. Clark, Blake Bordelon, Jacob A. Zavatone-Veth, Cengiz Pehlevan

## Abstract

Across many brain areas, neurons produce heterogeneous, seemingly disordered responses. Yet such circuits cannot be purely random, since they must possess some structure to generate the representations and computations underlying behavior. How much structure is present in recurrent connectivity relative to disorder, and how the interaction between the two shapes population dynamics and single-neuron responses, remain incompletely understood. Recurrent neural networks trained to perform tasks have become a leading model of such circuits, but conventional training yields a single point in a vast space of task-compatible solutions, with no systematic way to explore this space and no theory of how internal representations vary within it. Without such a theory, the questions above cannot be addressed, and comparisons between trained networks and neural data are difficult to interpret. Here, we introduce a control parameter that governs the degree to which learning reshapes recurrent connectivity, interpolating between a reservoir regime and one in which recurrent weights are restructured by learning to produce task-relevant internal representations. Varying this parameter generates a family of task-compatible solutions whose internal dynamics differ in a controlled and interpretable way. We derive a dynamical mean-field theory showing that, while population-level dynamics converge to a deterministic limit, individual neurons are driven by independent samples from a single-neuron input-current distribution. When connectivity is random, this distribution is Gaussian. Recurrent restructuring drives it toward task-dependent, non-Gaussian forms. In linear networks, restructuring amplifies task-relevant frequencies. In nonlinear networks, it drives a phase transition from chaotic, high-dimensional activity to ordered, low-dimensional dynamics that generalize temporally beyond the training period. We apply the theory to a reaching task in which a recurrent network must reproduce macaque muscle activity, and find that optimally matching simultaneous motor-cortex recordings requires only a small degree of restructuring, with learned structure coexisting with random heterogeneity. These results suggest a broader picture in which large recurrent circuits are largely random but contain, to varying degrees, structured recurrent connectivity sufficient for generalizable, task-relevant representations.^1^

## 1 Introduction

Neurons in early sensory areas often exhibit orderly response properties. In primary visual cortex, for example, individual neurons respond selectively to oriented edges [1], with the population tiling the space of orientations in an organized fashion [2]. Similarly orderly tuning has been observed in auditory and somatosensory cortices [3]. Inspired by these successes, early work in motor neuroscience sought analogous structure in motor cortical responses, seeming to find cells in primary motor cortex with cosine-like tuning to the direction of arm reaches [4].

Subsequent experiments and analyses indicated that, during reaching, motor cortical neurons produce multiphasic, heterogeneous, and highly dynamic responses that resist description via simple tuning curves [5, 6]. Such complexity appears to be the rule rather than the exception across large mammalian circuits. Head-direction cells, despite being organized around a one-dimensional variable, show striking response heterogeneity [7]. Hippocampal place fields are diverse and spatially irregular [8]. Neurons in prefrontal cortex carry mixed, condition-dependent signals during cognitive tasks [9]. Premotor neurons show richly diverse temporal responses during movement preparation and execution [10–12]. This pervasive heterogeneity may lead one to think that neural circuits are largely random, an idea with empirical support in systems such as the insect mushroom body and mammalian piriform cortex, where projections appear to be substantially unstructured [13–15].

However, random connectivity on its own cannot be the full story. Recurrent circuits with random connectivity, called *reservoir* networks, can generate feature selectivity [16] and perform pattern separation [17], but they generally produce high-dimensional chaotic activity [18, 19]. Such activity seems clearly at odds with the structured, low-dimensional population-level dynamics observed in motor cortex [6], in areas underlying cognitive tasks, and in other circuits [20]. Structured recurrent connectivity is also beneficial for generating representations that support generalization and compositionality [21, 22]; we analyze generalization in a simplified scenario below. Neural circuits thus likely contain both a degree of disorder in their connectivity, perhaps due to stochastic wiring rules [23], and some learned structure that generates task-relevant representations. How much structure is present relative to disorder, and how the interaction between them shapes population-level dynamics and single-neuron responses, remain incompletely understood.

Addressing these questions in a circuit model requires a way to endow the network with task-relevant connectivity. One approach, which has proven amenable to theoretical analysis, is to specify the desired neuronal representation and construct connectivity that supports it [24–26]. This allows one to study the resulting, generically disordered connectivity and network dynamics, but the neuronal responses themselves are fixed *a priori*. A more ambitious approach is task training, in which the representation is not imposed but learned, with the learning signal provided by asking the network to perform an experimentally relevant task. When optimized to perform motor, cognitive, or sensory tasks, such networks often develop neuronal responses and population-level dynamics that resemble those found in neural recordings [9– 11, 22, 27, 28], allowing such comparisons to inform our understanding of the circuit at hand.

Despite the ubiquity of task-trained RNNs in systems neuroscience, we lack a theory of the representations these networks learn. Conventional training yields a single point in a vast space of task-compatible solutions, with no systematic way to explore how representations vary along the order-disorder axis and no theory of what governs this variation. This has two consequences. First, the questions of structure and disorder posed above cannot be addressed in a principled way. Second, comparisons between trained RNNs and neural data are difficult to interpret. Ideally, one would have a controllable parameter that generates diverse task-compatible solutions, paired with a theory of how that parameter shapes internal representations, so that when a particular solution matches neural data one can draw specific conclusions about the underlying circuit (we return to this point in the Discussion). A related practical issue is that, without such a theory, it is unclear under what conditions population-level dynamics are consistent across training runs and network sizes.

Here, we develop a theoretical framework that addresses these issues. Drawing on modern machinelearning theory [29–34], we propose a parameterization for RNNs trained with backpropagation through time that guarantees consistent population-level behavior in large networks and, crucially, allows us to titrate the degree of structure and disorder in the learned connectivity. The titration is controlled by a single parameter *γ*. As *γ* → 0^+^, recurrent weights remain unstructured and only the readout is learned, yielding a reservoir regime [35, 36]. As *γ* increases, recurrent connectivity is progressively restructured by learning, producing task-dependent internal representations. Varying *γ* thus generates a continuous family of task-compatible solutions whose internal dynamics differ in a controlled and interpretable way.

To characterize this family, we develop a dynamical mean-field theory (DMFT) describing the network in the limit of a large number of neurons. The DMFT shows that the population-averaged correlation function is governed by an action whose minimum determines the network’s macroscopic behavior. The action contains two competing terms, one inherited from the classical theory of chaotic random networks [18] that favors unstructured connectivity and high-dimensional activity, and another arising from learning that pulls the network toward task-relevant, low-dimensional representations. The parameter *γ* controls the relative strength of these terms, and the learned network reflects the outcome of the competition. A key prediction is that population-level statistics converge to a deterministic limit, while individual neurons remain independent samples from a single-neuron response distribution whose form is also set by *γ*. In the reservoir regime, this distribution is Gaussian, reflecting random connectivity. Recurrent restructuring drives it toward task-dependent, non-Gaussian forms. At the level of weights, this restructuring pulls outlier eigenvalues out of the random bulk, introducing dynamical modes that carry the learned temporal structure. For linear networks, increasing *γ* reshapes the effective temporal filter, amplifying task-relevant frequencies. For nonlinear networks, it drives a phase transition from chaotic, high-dimensional activity to ordered, low-dimensional dynamics that support generalization to time points beyond the training window.

We apply this framework to a motor task, previously studied by Sussillo et al. [27], in which an RNN must reproduce electromyographic (EMG) signals recorded from eight muscles during macaque reaching. By varying *γ*, we obtain a family of networks that all reproduce the EMG targets with comparable accuracy but differ systematically in their internal dynamics. Comparing these networks to simultaneous motor-cortex recordings, we find that reservoir networks (*γ* → 0^+^) produce accurate muscle outputs but poorly match neural responses. An intermediate degree of recurrent restructuring substantially improves this match, though the strength of the improvement varies across metrics. The best-matching networks occupy a regime where learned dynamical structure coexists with random heterogeneity. More broadly, this picture suggests that large recurrent circuits may operate similarly, with varying degrees of structured recurrence embedded within otherwise random connectivity, sufficient to support generalizable, feature-based representations^2^.

## 2 Results

### 2.1 Recurrent-network model and training

We consider rate-based recurrent neural networks (RNNs) of *N* neurons. Each neuron *i* = 1, …, *N* has a preactivation *x*_*i*_(*t*) and an activation *ϕ*_*i*_(*t*) ≡ *ϕ*(*x*_*i*_(*t*)), where *ϕ*(·) is a pointwise nonlinearity. As is typical, we interpret *x*_*i*_(*t*) as a filtered input current to neuron *i* and *ϕ*_*i*_(*t*) as its firing rate. Except when otherwise specified, we use *ϕ*(·) = tanh(·) throughout this work.

The neurons are coupled through a matrix of recurrent weights *J*_*ij*_ and receive *D*_in_-dimensional time-dependent inputs *I*_*a*_(*t*), where *a* = 1, …, *D*_in_, via input weights *U*_*ia*_. The network dynamics follow

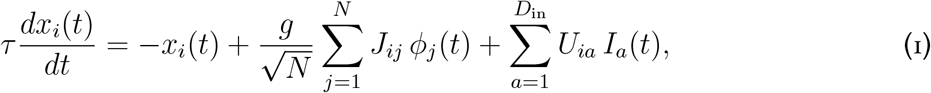

where *τ* is a single-neuron time constant that, until Sec. 2.6, we set to unity. The factor 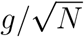 ensures that the typical size of the recurrent input to each neuron remains fixed as *N* grows, and the gain parameter *g* controls the strength of recurrent interactions [18]. The network produces *D*_out_-dimensional outputs *y*_*a*_(*t*), where *a* = 1, …, *D*_out_, through a linear readout with weights *V*_*ia*_,

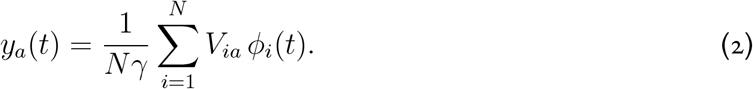

The parameter *γ >* 0 plays a central role in what follows, serving as our primary knob for titrating the degree of recurrent restructuring. A scaling argument explains why. The activations *ϕ*_*i*_(*t*) and the readout weights *V*_*ia*_ are both 𝒪(1) in *N*. When the activations have generic alignment with the columns of the readout weight matrix, as is the case at initialization and in the absence of recurrent restructuring, the sum 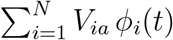 grows as 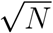. After the 1*/*(*Nγ*) prefactor in the readout, the output therefore shrinks as 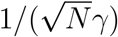, vanishing as *N* → ∞ for any fixed *γ*. To produce a finite output, the activations *ϕ*_*i*_(*t*) must develop alignment with the columns of *V*_*ia*_, requiring the internal representation to adapt to the task. Larger *γ* demands stronger alignment and thus more restructuring of the recurrent weights. Smaller *γ* allows the 1*/*(*Nγ*) prefactor to compensate, so that even weak alignment suffices. In the limit *γ* → 0^+^, the recurrent weights are not adapted by learning at all, and the network operates as a reservoir [35, 36, 38], with only the readout weights effectively trained (Fig. 1A). In the machine-learning theory community, this phenomenon is known as *feature learning* [34], with the small-*γ* and large-*γ* limits referred to as the *lazy* and *rich* regimes, respectively [29]. The parameter scalings we have adopted correspond to the so-called *maximal-update parameterization* (*µ*P) [30–32, 34], which ensures that the degree of feature learning remains nontrivial and consistent as *N* grows.

**Figure 1:**
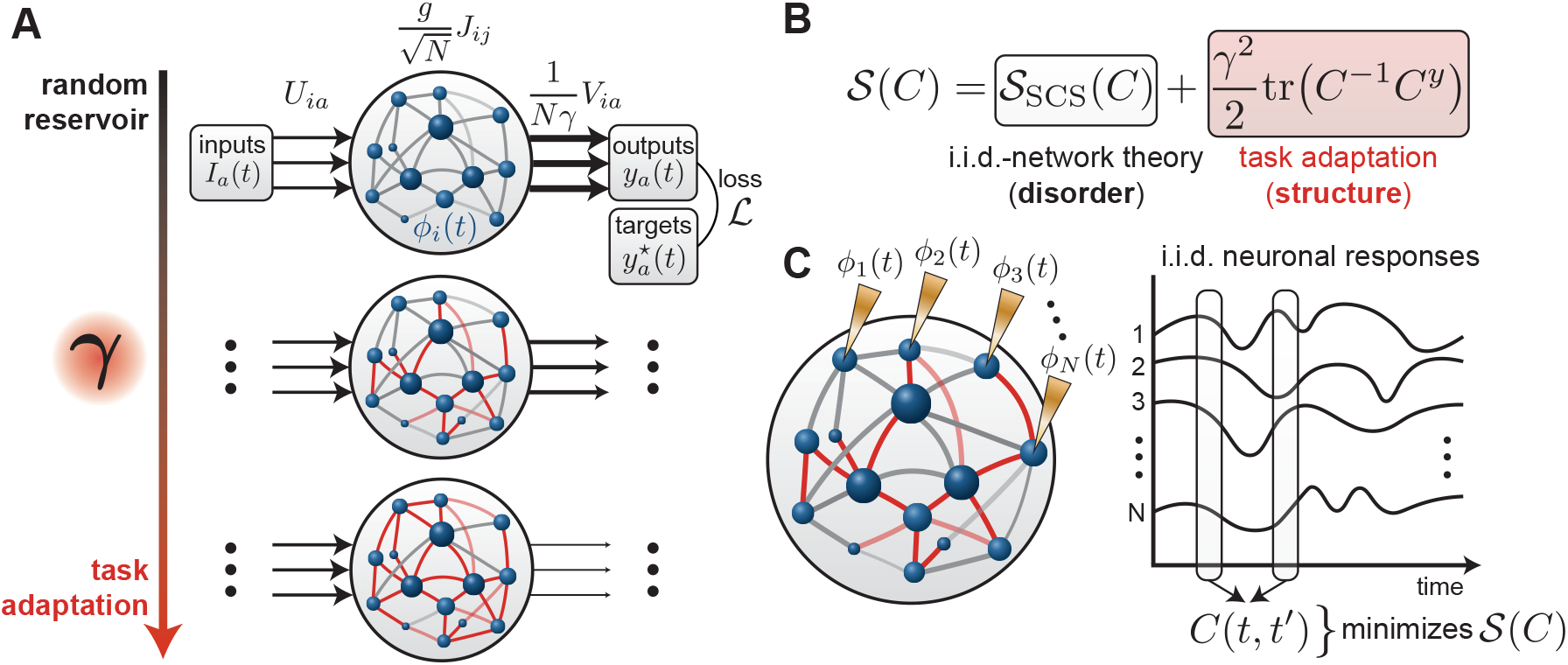
Framework overview. **(A)** Illustration of our task-trained RNN framework. An RNN receives time-dependent inputs, processes them through recurrent connectivity, and produces outputs through a linear readout. The network is trained to match target outputs by minimizing a loss. The parameter *γ* controls the degree to which learning reshapes recurrent connectivity, interpolating between a reservoir regime (*γ* → 0^+^, top) in which recurrent weights remain unstructured, and a task-adapted regime (large *γ*, bottom) in which learned structure progressively reshapes the recurrent connections. In machine learning theory, these are referred to as the lazy and rich regimes, respectively [29, 30]. **(B)** In the large-network limit, the population-averaged correlation function *C*(*t, t*′) is deterministic and minimizes an action 𝒮 (*C*) composed of two competing terms. The first is inherited from the mean-field theory of random networks and favors chaotic, high-dimensional activity. The second arises from learning and drives the network toward temporal structure aligned with the task targets. **(C)** Individual neurons in the large-network limit are independent and identically distributed (i.i.d.) samples from a distribution over response profiles. That is, each neuron produces a temporal trace that is a random draw from a distribution, with the form of this distribution depending on the degree of recurrent restructuring.

We train the network in an end-to-end manner, meaning that the recurrent weights ***J***, input weights ***U***, and readout weights ***V***, collected in the parameter set **Θ**, are all learned jointly. The training objective is to make the network output *y*_*a*_(*t*) match a target sequence 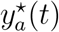 over a time interval [0, *T*]. We define a loss function, interpretable within our theoretical framework as an energy, consisting of a mean-squared-error (MSE) data term and a quadratic or ridge regularizer,

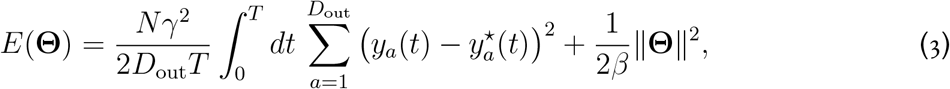

where ∥**Θ**∥^2^ is the sum of squares of all parameters and the prefactor *Nγ*^2^ ensures compatibility between the data term and the output scaling in Eq. (2). The output *y*_*a*_(*t*) depends on the parameters **Θ** through the recurrent dynamics of Eqs. (1)–(2), making this a highly nonlinear optimization problem. The scalar *β >* 0 controls both the strength of the ridge regularization and the amplitude of the learning noise introduced below. The factor of 1*/β* in the ridge term ensures that even in the limit *β* → ∞, where the noise vanishes and the data-fit term dominates, the regularization also vanishes, so that the learning dynamics sample a degenerate manifold of solutions rather than converging to a single point [39]. The formulation extends straightforwardly to multiple input-output sequence pairs by averaging the data term over a set of sequences (see Sec. 2.6).

We learn the parameters through a continuous-time Langevin dynamics in which the parameters **Θ** evolve over a learning time *s*,

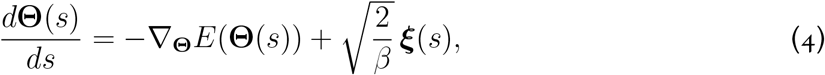

where ***ξ***(*s*) is white noise, independent across parameters and learning time. The first term is the gradient of the energy, computed via backpropagation through time, and drives the parameters toward configurations that fit the target while keeping weights small. The second term injects noise whose amplitude is controlled by *β*, so that larger *β* corresponds to less noise and a stronger emphasis on minimizing the energy. We refer to this as Langevin gradient flow. A key property is that as *s* → ∞, the distribution over parameters converges to the Gibbs distribution, *P* (**Θ**) ∝ exp (−*β E*(**Θ**)). At large *β*, this distribution concentrates on low-energy configurations, that is, networks that fit the target well while keeping weights moderate. Working with the Gibbs distribution allows us to characterize the statistical properties of trained networks at equilibrium without tracking the full learning trajectory.

The Gibbs distribution has a natural Bayesian interpretation as a posterior that balances an independent and identically distributed (i.i.d.) Gaussian prior on the weights against a likelihood that rewards alignment between the network’s temporal correlations and those of the target. The prior favors random, unstructured connectivity; the likelihood favors task-relevant structure. The parameter *γ* controls how far the posterior departs from the prior, titrating the mixture of randomness and task structure in the learned connectivity [34, 39–41].

### 2.2 Large-network limit and dynamical mean-field theory

In the large-*N* limit, the network can be analyzed using dynamical mean-field theory (DMFT). The core idea is to reduce the *N*-dimensional dynamics to a small number of self-consistent equations for order parameters (summary statistics) that describe the population-level behavior. These order parameters are self-averaging, meaning that they can be measured in finite networks and agree with the DMFT predictions up to fluctuations that shrink as 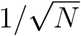, independent of the particular weight realization [42]. DMFT therefore provides an exact description of the population-level dynamics as *N* → ∞.

The central order parameter is the population-averaged temporal correlation function,

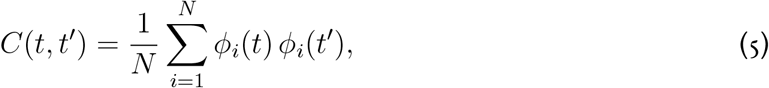

which measures the similarity of the network’s internal representation between times *t* and *t*′. For a random network with i.i.d. weights and no learning, the DMFT was developed by Sompolinsky, Crisanti, and Sommers (SCS) [18]. One way to formulate their theory, which will prove natural for incorporating learning, is in terms of an action 𝒮_SCS_(*C*) that *C*(*t, t*′) minimizes. The action plays a role analogous to a loss function, but for the population-level structure of the network rather than its parameters; the physical correlation function is the one that minimizes it. The SCS action encodes the statistics of a single neuron driven by a self-consistent Gaussian process whose covariance depends on *C*(*t, t*′) itself (Sec. SI.1). Our theory shows that learning adds a second term to this action. In the *β* → ∞ limit, the correlation function of the learned network minimizes

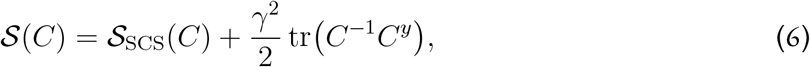

where 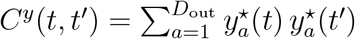 is the temporal correlation of the target outputs, and the trace and inverse in the second term treat *C*(*t, t*′) and *C*^*y*^(*t, t*′) as matrices with temporal indices. The structure of this action makes the competition between randomness and task structure explicit (Fig. 1B). The first term, inherited from the classical SCS theory, encodes the statistics of a random, unstructured network. The second term penalizes misalignment between the network’s temporal correlations and the target. When *C*(*t, t*′) has little power along temporal modes present in *C*^*y*^(*t, t*′), the trace tr(*C*^−1^*C*^*y*^) is large, driving *C*(*t, t*′) to better capture those modes. The strength of this drive is set by *γ*^2^, so as *γ* → 0^+^ the action reduces to 𝒮_SCS_(*C*) and the network behaves as a reservoir, while at large *γ* the learned term dominates and the correlation structure is shaped heavily by the task. In the *β* → ∞ limit, the network output matches the target exactly, 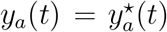, provided *C*(*t, t*′) is invertible (Eq. (SI.13)). At finite *β*, the inverse *C*^−1^(*t, t*′) in the action is replaced by a regularized inverse, and the output incurs a computable discrepancy from the target (Sec. SI.1).

### 2.3 Learning reshapes the single-neuron response distribution

At large *N*, while the correlation function *C*(*t, t*′) is deterministic, individual neurons are independent, identically distributed samples from a distribution over response profiles (Fig. 1C). Neurons are thus heterogeneous at all values of *γ*, but the character of this heterogeneity changes with *γ*, reflecting the mixture of randomness and task structure in the underlying connectivity. We now describe this distribution by unpacking the structure of the action.

The SCS action 𝒮_SCS_(*C*) contains an inner optimization over a conjugate (or auxiliary) variable *Ĉ*(*t, t*′), so that the full variational problem is a saddle point (Sec. SI.1). A saddle point is a stationary point of the action (where all gradients vanish) that is a minimum in some directions and a maximum in others; here, the action is minimized over *C*(*t, t*′) and maximized over *Ĉ*(*t, t*′). The stationarity condition with respect to *Ĉ*(*t, t*′) produces a self-consistency equation that determines *C*(*t, t*′), while the saddle-point value of *Ĉ*(*t, t*′) itself controls how the distribution over single-neuron responses departs from a baseline Gaussian. In the SCS DMFT, each neuron receives an input current *η*(*t*) drawn independently from a Gaussian process with mean zero and covariance ⟨*η*(*t*) *η*(*t*′)⟩ = *g*^2^*C*(*t, t*′) + *C*^*I*^(*t, t*′), where *C*^*I*^(*t, t*′) is the covariance of the external inputs. This input current replaces the actual recurrent input from other neurons, capturing its statistics in a self-consistent way. The current is passed through the single-neuron dynamics (1 + ∂_*t*_) *x*(*t*) = *η*(*t*) to produce the preactivation *x*(*t*), which, because this is a linear filter of a Gaussian process, is also Gaussian. Applying the nonlinearity then gives the activation *ϕ*(*t*) = *ϕ*(*x*(*t*)). In the learned system, a crucial step is inserted into this process, namely, the Gaussian distribution over input currents is reweighted by a tilting factor 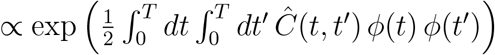, where *ϕ*(*t*) is the activation produced by *η*(*t*) through the single-neuron dynamics. This tilting biases the population toward neurons whose input currents produce activations aligned with the structure encoded in *Ĉ*(*t, t*′). Self-consistency requires that the correlation function computed under this tilted distribution matches the order parameter *C*(*t, t*′) that enters the action.

As *γ* → 0^+^, the learning term in the action (6) vanishes and the saddle-point value of the conjugate variable is *Ĉ*(*t, t*′) = 0. The tilting factor reduces to a constant, the input current *η*(*t*) remains a Gaussian process, and the SCS theory is recovered. When *γ >* 0, the learning term shifts the saddle-point value of *C*(*t, t*′) away from the random-network solution, which in turn drives *Ĉ*(*t, t*′) away from zero. The resulting tilting makes the distribution over input currents non-Gaussian, driving the preactivation and activation distributions away from Gaussianity in manner that depends on the task through *C*^*y*^(*t, t*′). Thus, although neurons are always i.i.d. samples from a population distribution, the distribution itself is progressively reshaped by learning. As *γ* → 0^+^ it is Gaussian, reflecting purely random connectivity, while at larger *γ* it acquires non-Gaussian, task-relevant structure. Intuitively, recurrent restructuring enriches the population with neurons whose temporal responses are useful for the task, at the expense of those that are not. The tilting factor implements this enrichment, exponentially upweighting input current realizations whose resulting activations have two-point structure aligned with *Ĉ*(*t, t*′), which the saddle-point conditions tie to the task target *C*^*y*^(*t, t*′). Non-Gaussianity in single-neuron statistics is the imprint of this task-driven enrichment.

### 2.4 Recurrent restructuring amplifies task-relevant frequencies in linear networks

As *γ* → 0^+^ (the SCS theory), the distribution over single-neuron responses is Gaussian, and the self-consistency equations for the correlation function can be solved analytically. At *γ >* 0, the non-Gaussian tilting introduced by learning necessitates numerical solutions (Sec. SI.1). To build analytical intuition for how recurrent restructuring reshapes activity, we consider the one case in which a closed-form solution can still be obtained, namely linear recurrent networks, with *ϕ*(·) the identity. Linear networks are limited as models of biological circuits and cannot exhibit phenomena such as chaos, but afford a tractable testbed. The dynamics are linear, but the learning process that produces the trained weights is not [34, 43, 44], so obtaining exact results for the learned representations is nontrivial.

A further simplification arises if we assume that the inputs, outputs, and network activity are statistically stationary, meaning that correlation functions depend only on time differences. Under this assumption, the DMFT equation for the power spectrum *C*(*ω*) decouples across frequencies, reducing to a quadratic equation at each *ω* in the *β* → ∞ limit (Sec. SI.3.3). This yields a closed-form expression for the power spectrum of the learned network in terms of the input and target power spectra,

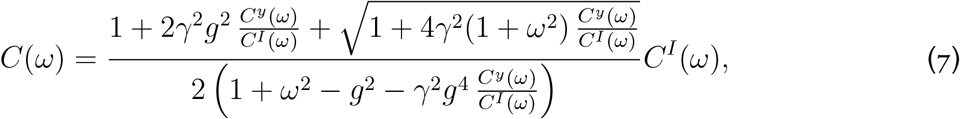

where *C*^*I*^(*ω*) and *C*^*y*^(*ω*) are the power spectra of the input and target, respectively. The ratio *C*^*y*^(*ω*)*/C*^*I*^(*ω*) measures how much each frequency is over-represented in the target relative to the input, and learning reshapes the network’s transfer function to amplify precisely those frequencies.

We first examine the reservoir limit (*γ* → 0^+^), in which Eq. (7) reduces to *C*(*ω*) = *C*^*I*^(*ω*)*/*(1 + *ω*^2^ − *g*^2^). For white-noise input (*C*^*I*^(*ω*) = 1; Fig. 2A), this is the power spectrum of an Ornstein-Uhlenbeck process with correlation time 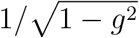, which diverges as *g* → 1^−^, reflecting the approach to instability of the random linear network. The reservoir power spectrum decays monotonically from low to high frequencies (Fig. 2B, *γ* → 0^+^ curves in each panel).

**Figure 2:**
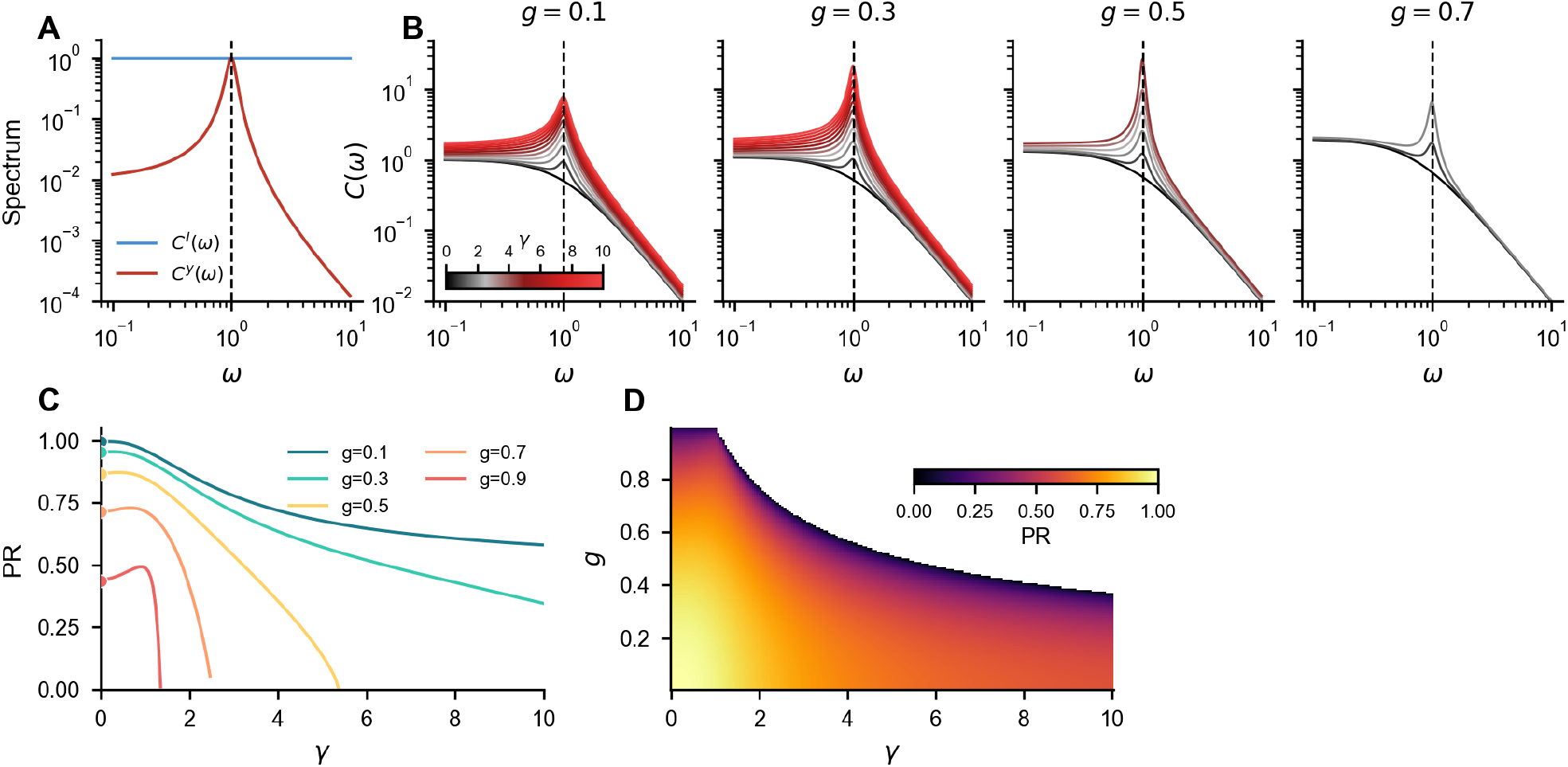
Recurrent restructuring amplifies task-relevant frequencies in linear networks. **(A)** Linear networks are trained to transform white-noise input with flat power spectrum *C*^*I*^(*ω*) = 1 into outputs whose power spectrum is concentrated around a preferred frequency *ω*_⋆_ = 1 (dashed vertical line). **(B)** Power spectra *C*(*ω*) of the learned network activity for different values of the gain *g* (panels). Within each panel, increasing *γ* (indicated by hue) progressively amplifies power near the target frequency. For a given *g*, there is a maximum *γ* beyond which the linear dynamics become unstable (Sec. SI.3.3); only stable parameter combinations are shown. **(C)** Participation ratio of the stationary covariance (defined in main text) as a function of *γ* for several values of *g*. **(D)** PR over the (*γ, g*) plane. The blank region corresponds to unstable linear dynamics.

We now consider the effect of increasing *γ*. We illustrate with a network trained to transform whitenoise input into noise whose power spectrum is sharply peaked around a preferred frequency *ω*_⋆_ = 1, with target spectrum *C*^*y*^(*ω*) = 1*/*(1 + 10^2^(|*ω*| − *ω*_⋆_)^2^) (Fig. 2A). As *γ* increases from zero, a peak emerges around the target frequency and grows progressively (Fig. 2B). Recurrent restructuring thus selectively amplifies the frequencies demanded by the task. Equation (7) also implies a stability condition, namely that the denominator must remain positive for all *ω*, which places an upper bound on *γ* for a given *g* and target spectrum. For each value of *g*, we show power spectra only for values of *γ* at which the linear dynamics remain stable.

We summarize the effect of recurrent restructuring on the population-level dynamics using the participation ratio (PR) of the power spectrum, 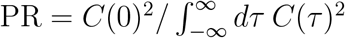, which measures the number of dimensions explored by the network per unit of time [45]. In the reservoir limit, 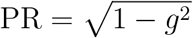, which vanishes as *g* → 1^−^, consistent with the diverging correlation time noted above [46]. With increasing *γ*, the participation ratio first increases, prominently so for large *g*, and then decreases toward zero as the linear dynamics approach the stability boundary (Fig. 2C,D). This nonmonotonic dependence of dimensionality on *γ* will reappear in the nonlinear networks studied below.

A linear network can only output frequencies already present in its input and cannot autonomously generate temporal structure. We turn next to nonlinear networks, where richer dynamical phenomena become possible.

### 2.5 Recurrent restructuring suppresses chaos and enables temporal generalization

A hallmark of task-adapted representations in feedforward networks is that they support generalization [30, 47–52]. We now ask whether an analogous phenomenon occurs in the temporal domain, that is, whether restructuring recurrent connectivity enables generalization across time. Unlike the linear networks analyzed above, nonlinear networks can generate novel temporal structure autonomously, and we therefore turn to a common task in the RNN literature that probes exactly this question, the autonomous generation of a periodic signal [28, 53]. Producing a stable oscillation requires the suppression of chaotic dynamics, making this task a natural setting for understanding how recurrent restructuring reshapes the dynamical regime of the network.

We trained networks to produce a single period of a two-dimensional sinusoidal output with period *T* = 10 time constants, with no external input (Fig. 3A). The target returns to its initial value at time *T*, which is compatible with periodicity but does not imply it, since the network sees only one period and has no opportunity to learn from repetitions. After training, we ran the network beyond the training window and examined its generalization behavior. We restrict to the regime *g >* 1, in which the dynamics without recurrent restructuring are chaotic, as is standard in practice when training RNNs [53].

**Figure 3:**
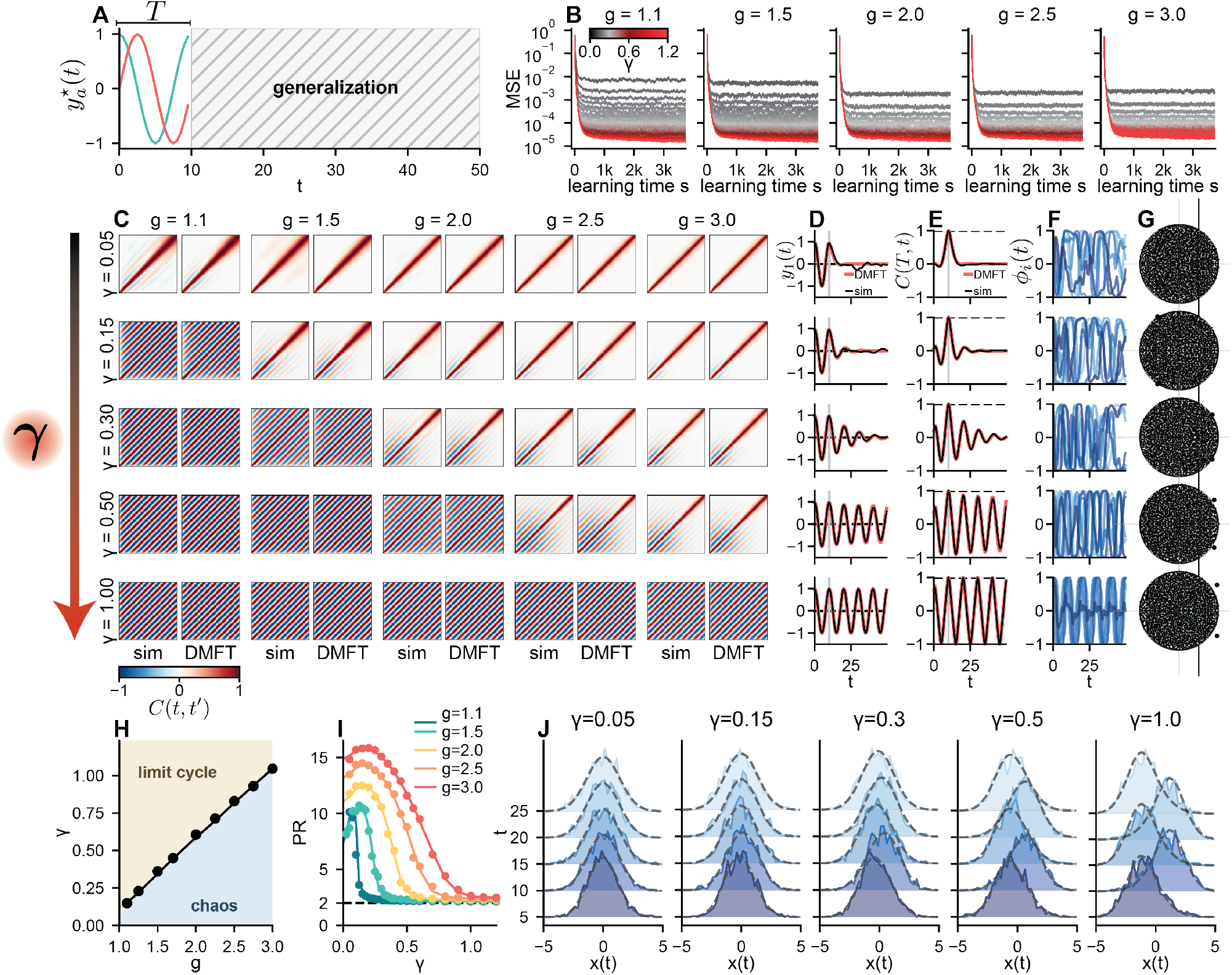
Recurrent restructuring suppresses chaos and enables temporal generalization. **(A)** Target signal 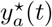: a single period (*T* = 10) of a two-dimensional sinusoidal output. We asked what dynamics trained networks produced at later times, *T < t* ≤ *T*_tot_, where *T*_tot_ = 50 (generalization window). **(B)** For each pair (*g, γ*), we trained 10 independent networks of size *N* = 2500 via Langevin gradient flow (*β* = 2000). Here we show learning curves for different values of *γ* (lines) and *g* (columns). All networks achieved low training error, with smaller residuals at larger *γ* (a finite-*β* effect; see main text). **(C)** Temporal correlation *C*(*t, t*′), over the *T*_tot_ × *T*_tot_ window, for different values of *γ* (rows) and *g* (columns), comparing simulation (left in each panel, averaged across 10 networks) and DMFT (right in each panel). **(D)** Network output *y*_1_(*t*) across the training and test periods, for *g* = 2 and the same *γ* values as in (C). Solid lines show the output of a single trained network; dashed lines show the DMFT prediction obtained by kernel regression from the training-window correlation function (Eq. (SI.33)). **(E)** Slices *C*(*T, t*) of the temporal correlation for *g* = 2 and the same *γ* values as in (C), comparing simulation (thin lines) and DMFT (thick lines). Vertical line indicates *t* = *T*. **(F)** Single-neuron activation traces across the training and test periods for *g* = 2 and the same *γ* values as in (C). **(G)** Eigenvalue spectra of 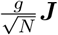 for *g* = 2 and the same *γ* values as in (C). Black dots show eigenvalues from a single network, and gray dots show eigenvalues from the remaining 9 networks. At small *γ*, eigenvalues follow the circular law with radius *g*. Increasing *γ* pulls complex-conjugate outlier pairs out of the bulk. The vertical line indicates the stability boundary, Re(*λ*) = 1. **(H)** Phase diagram in the (*g, γ*) plane, with the curve *γ*^⋆^(*g*) separating the chaotic regime (large *g*, small *γ*) from the limit-cycle regime (small *g*, large *γ*); shaded regions are labeled accordingly. Dots show simulation, line shows DMFT. **(I)** Participation ratio of *C*(*t, t*′) over the training and test periods as a function of *γ* for different values of *g*, comparing simulation (dots, computed from the mean correlation across 10 networks) and DMFT (lines). **(J)** Preactivation distributions at selected time points for *g* = 2 and the same *γ* values as in (C), comparing simulation (filled histograms, 100 values collected from each of the 10 networks per *γ*) and DMFT (dashed outlines, includes tilting).

We trained networks and solved the DMFT equations numerically across a grid of (*g, γ*) values (Sec. SI.2, SI.5). All configurations achieve low training error (Fig. 3B), with smaller errors at larger *γ*. Any nonzero error is a finite-*β* effect, since the *β* → ∞ readout reproduces the target exactly whenever *C*(*t, t*′) is invertible (Eq. (SI.13)). To leading order in 1*/β*, the residual is proportional to *C*^−1^(*t, t*′)*Y* ^⋆^, with squared norm (*Y* ^⋆^)^⊤^*C*^−2^(*t, t*′)*Y* ^⋆^. The action’s learning term penalizes (*Y* ^⋆^)^⊤^*C*^−1^(*t, t*′)*Y* ^⋆^ with prefactor *γ*^2^, the same quadratic form in *Y* ^⋆^ but with *C*^−1^ in place of *C*^−2^. Larger *γ* shrinks both, and the residual along with them.

Despite comparable training performance, networks at different *γ* differ markedly in their internal dynamics and generalization behavior. Fig. 3C displays the correlation function *C*(*t, t*′) for each (*g, γ*) configuration, computed both from finite-*N* simulations and from the DMFT. The DMFT provides an extremely close match throughout, as confirmed by the overlaid slices *C*(*T, t*) in Fig. 3E. Beyond population-level statistics, the DMFT also predicts the network’s autonomous output in the test window via a kernel regression on the training-window correlation function (Eq. (SI.33), Sec. SI.2.2), and the prediction matches simulation closely throughout (Fig. 3D). At small *γ, C*(*t, t*′) decays rapidly away from the diagonal, the signature of chaotic dynamics with short-lived temporal correlations. As *γ* increases, oscillatory structure emerges and grows until, at large *γ*, it dominates and the chaotic decay is no longer visible.

Single-neuron activity traces reflect this same progression (Fig. 3F). Recall that individual neurons are i.i.d. samples from a distribution that is progressively reshaped by recurrent restructuring. At small *γ*, each neuron’s response is a smooth but random temporal fluctuation, and the population looks disordered. At large *γ*, responses converge toward sinusoids passed through the nonlinearity *ϕ*(·), differing only in phase, and the appearance of disorder is lost.

This progression culminates in a transition between chaotic and ordered dynamics. Following [54], we identified the onset of nonchaotic, limit-cycle dynamics by checking whether the normalized correlation function returned to unity after a time lag, indicating that the population vector of activity returns exactly to itself. We assessed this after 3*T* of autonomous running to allow convergence to a putative limit cycle. This criterion defines a critical value *γ*^⋆^(*g*) above which the dynamics are nonchaotic, partitioning the (*g, γ*) plane into chaotic and limit-cycle regimes (Fig. 3H). The DMFT predicts this phase boundary accurately. Larger *g* requires larger *γ* to suppress chaos, as stronger recurrent gain produces more vigorous chaotic fluctuations that recurrent restructuring must overcome.

In the non-stationary setting considered here, we compute a participation ratio over a finite observation window, 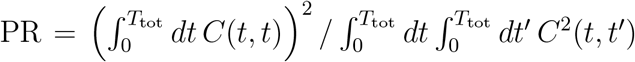, which counts the number of dimensions explored by network activity over this window, where *T*_tot_ = 50 (Fig. 3I). Unlike the stationary PR of Sec. 2.4, which measures a rate, this is a dimensionless count; the two correspond to different orders of limits in *N* and the observation time *T*_tot_ (Sec. SI.4). As *γ* increases from zero, PR first rises, then falls, eventually approaching two, the value expected for a pure sinusoidal oscillation. This nonmonotonicity arises because at small *γ* all variance comes from high-dimensional chaos, at large *γ* it comes from the two-dimensional oscillation, and at intermediate *γ* both sources contribute. This is qualitatively reminiscent of the nonmonotonic participation ratio observed in the linear case (Fig. 2C), despite the different definitions.

Fig. 3J shows the distributions of preactivations across RNN neurons at selected time points, comparing simulations with the DMFT. At small *γ*, distributions are Gaussian and centered at zero at all times, as expected from the SCS theory. As *γ* increases, the distributions develop mild skewness, and the peak oscillates back and forth over time rather than remaining centered at zero, both of which reflect the tilting mechanism of the DMFT.

The quantities examined so far (namely, the correlation function, neuronal traces, and preactivation distributions) are all accessible through the DMFT. To understand the recurrent restructuring underlying these changes at the level of the weights, we examined the eigenvalue spectra of the recurrent weight matrix 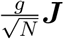 (Fig. 3G). As *γ* → 0^+^, eigenvalues are uniformly distributed within a disk of radius *g*, following the circular law for i.i.d. random matrices. Increasing *γ* pulls a pair of complex-conjugate outlier eigenvalues out of the bulk, with their distance from the bulk growing with *γ*, corresponding to progressively stronger oscillatory dynamics. This spectral structure provides a complementary view of the competition between randomness and task structure that the DMFT formalizes. In particular, the random bulk reflects the disordered component of single-neuron responses, while the outlier eigenvalues carry the learned oscillatory dynamics required for the task.

#### Frequency amplification, nonlinearly revisited

With the nonlinear DMFT in hand, we can revisit the frequency-amplification analysis of Sec. 2.4, now asking a *nonlinear* network, in the time-translation-invariant setting, to *autonomously* generate a target whose power is concentrated at frequency *ω*_⋆_ (rather than asking a linear network to produce such a target in response to a white-noise input; Sec. SI.3.4). We take the target to be a pure sinusoid. This target would be problematic for a linear network, with singularities produced in the transfer function (Eq. (7)). The nonlinearity resolves this because application of *ϕ*(·) in the time domain globally reshapes the power spectrum, producing continuous spectral content (with peaks at odd harmonics of *ω*_⋆_; Fig. SI.2). The nonlinearity also enables a qualitatively new phenomenon. In a linear network, frequencies are not mixed; the activity at any given frequency depends only on the input at that same frequency. Learning therefore has only one job, namely to amplify the frequencies that appear in the target, as seen in the power spectra of Fig. 2B. There is nothing to suppress, because the linear activation cannot generate power at frequencies absent from the input. The nonlinearity changes this. By acting pointwise in time, *ϕ*(·) produces harmonics and otherwise spreads power across frequencies, so the learning term must do double duty, both amplifying task-relevant frequencies and *suppressing* task-irrelevant ones. The DMFT solution makes this manifest. The conjugate variable *Ĉ*(*ω*), which in the linear theory is non-negative everywhere (Eq. (SI.49)), now becomes positive near *ω*_⋆_, where it amplifies, and broadly negative elsewhere, where it suppresses (Fig. SI.2). The harmonics produced by the nonlinearity itself fall within this suppressed region and are pruned along with the rest of the task-irrelevant spectral content.

### 2.6 A family of networks that solve a macaque reaching task

The preceding sections illustrated our framework in simplified settings, with frequency amplification in linear networks and suppression of chaos with temporal generalization in nonlinear networks. In both cases, recurrent restructuring introduced task-relevant structure into an otherwise random network, pulling outlier eigenvalues out of the random spectral bulk, reshaping population-level activity, and driving single-neuron statistics away from Gaussianity. We now apply this framework to a realistic motor task and show that these same phenomena arise. In the next section, we ask which degree of recurrent restructuring best accounts for neural data.

The task is drawn from Churchland et al. [6], in which a macaque performed 27 reaching movements, including both direct reaches and curved reaches around obstacles, producing diverse hand trajectories (Fig. 4A, inset) and muscle activation patterns. Following Sussillo et al. [27], who originally studied this task in the task-trained RNN framework, we trained an RNN to transform temporally simple inputs into the temporally complex EMG patterns recorded from 8 muscles during these reaches. The network receives two types of input (Fig. 4A, left). The first is a six-dimensional condition-specific signal derived from the top principal components of the recorded preparatory neural activity, which specifies which of the 27 reaches to perform. The second is a condition-independent hold cue whose offset precedes movement.

**Figure 4:**
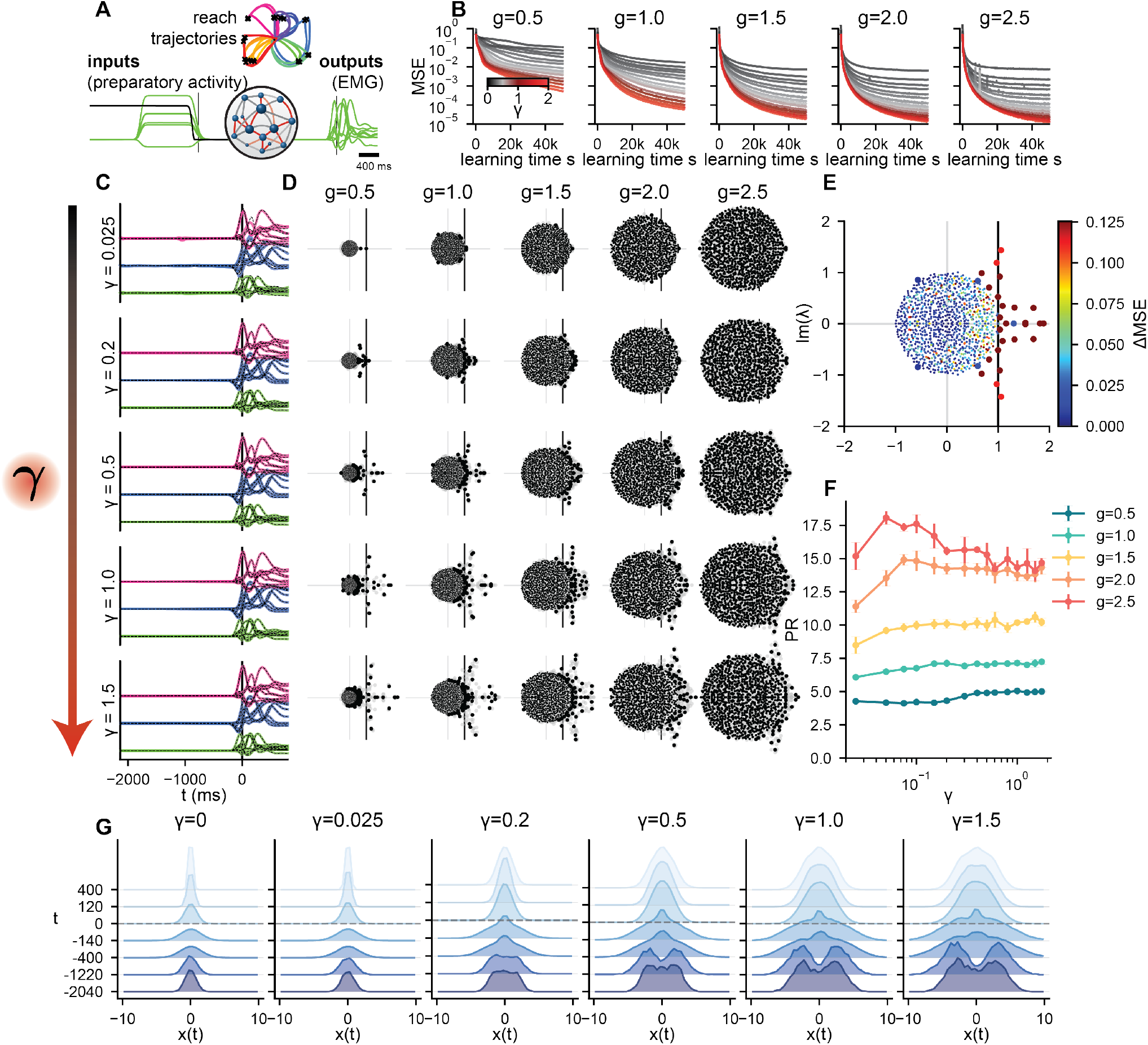
A family of networks that solve a macaque reaching task. **(A)** Task schematic. The network receives a six-dimensional condition-specific input derived from preparatory neural activity and a condition-independent hold cue (left), and produces eight-muscle EMG output (right). Inset: hand trajectories for all 27 reach conditions. **(B)** For each pair (*g, γ*), we trained 5 independent networks of size *N* = 1024 via Langevin gradient flow (*β* = 10^6^). Here we show the MSE as a function of learning time *s* for different values of *γ* (lines) and *g* (columns), averaged across 5 networks per condition. **(C)** EMG traces for *g* = 1.5 and various values of *γ* (rows), showing three representative reach conditions (colors as in (A) inset). Translucent colored lines show the trained RNN output; thin dashed black lines show the target (measured) EMG. The vertical line indicates movement onset (*t* = 0). **(D)** Eigenvalue spectra of 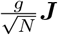 for the same *γ* values as in (C) (rows) and different values of *g* (columns). Black dots show eigenvalues from a single network, and gray dots show eigenvalues from the remaining networks. The vertical line indicates the stability boundary, Re(*λ*) = 1. **(E)** Eigenvalue spectrum for *g* = 1 and *γ* = 0.3, with each eigenvalue colored by the increase in MSE (ΔMSE) when the corresponding mode was ablated. **(F)** Participation ratio of RNN activity computed over a ± 400 ms window around movement onset, as a function of *γ* for different values of *g* (colors). **(G)** Preactivation distributions at selected time points for *g* = 1 and the same *γ* values as in (C), plus an untrained network (*γ* → 0^+^).

We study monkey J, as analyzed in the main text of Sussillo et al. [27]; simultaneous neural recordings from M1 and PMd are from Churchland et al. [6] and will be used for comparison in the next section. We set the time constant to *τ* = 50 ms and trained networks of *N* = 1024 neurons across a grid of *g* and *γ* values, with 5 independent runs per configuration via Langevin gradient flow (Sec. SI.2). Task parameters were matched to Sussillo et al. [27] wherever possible (Table SI.2; Sec. SI.6). For this task, the DMFT equations cannot currently be solved numerically because of issues of computational complexity and stability, an important avenue for future work. The theory nonetheless guarantees existence of a large-*N* limit, provides expectations for the role of *γ*, and identifies which summary statistics are constrained by the task [55].

All configurations produce accurate EMG output (Fig. 4C), with the MSE low throughout the (*g, γ*) grid (Fig. 4B), though somewhat higher at small *g* and small *γ*. Even near the reservoir limit, the network closely tracks the target muscle activity across conditions. By varying *γ*, we thus obtain a family of networks that all solve the same task but differ systematically in their internal dynamics, providing a controlled way to navigate the space of task-compatible solutions. Sussillo et al. [27] explored a version of this idea by comparing two networks trained with different regularization schemes, one that matched neural data well and one that did not, but without the ability to continuously modulate the nature of recurrent restructuring, and without a theory linking such restructuring to internal dynamics. Our framework places this comparison on systematic footing, generating a continuous family whose internal representations vary in an interpretable way.

We examined the eigenvalue spectra of the recurrent weight matrix (Fig. 4D). As in the sine wave task (Fig. 3G), outlier eigenvalues emerge from the random bulk with increasing *γ*, but here the spectral structure is richer, reflecting the higher-dimensional nature of the task. Several outliers occur in complex-conjugate pairs with large imaginary parts, corresponding to oscillatory modes consistent with the quasi-oscillatory dynamics identified in motor cortex by Churchland et al. [6]. Sussillo et al. [27] observed similar spectral structure by linearizing their regularized model around its fixed point. We confirmed the functional relevance of these outliers by ablating individual eigenvalues and measuring the resulting change in MSE (Fig. 4E). Removing outlier eigenvalues, particularly those with large imaginary parts, substantially degrades task performance, while removing bulk eigenvalues has negligible effect.

We computed the participation ratio of *C*(*t, t*′) over a ±400 ms window around movement onset as a measure of effective dimensionality (Fig. 4F). As in the linear (Fig. 2C) and sine wave (Fig. 3I) cases, PR is nonmonotonic for large *g*, first rising as task-relevant modes are added to the high-dimensional chaotic activity, then falling as those modes come to dominate. This nonmonotonicity is prominent at large *g* because it is in this regime that the inputs fail to suppress chaotic activity, at least at small *γ*, so that both chaotic and task-relevant variance coexist.

Fig. 4G shows the distributions of preactivations across RNN neurons at representative time points. As *γ* → 0^+^, preactivations are approximately Gaussian, as expected from random connectivity, with slight positive excess kurtosis reflecting temporal structure in the inputs. Increasing *γ* reshapes these distributions into non-Gaussian forms whose character depends on the phase of the task. Early in the trial, the distribution is bimodal. Around movement onset, it develops a sharp central peak. Later in the movement, it returns to a roughly Gaussian shape. These time-varying departures from Gaussianity are a manifestation of the tilting mechanism of Sec. 2.3.

The phenomena identified in simplified settings, namely outlier eigenvalues emerging from the random bulk, non-Gaussian single-neuron statistics, and nonmonotonic dimensionality, all carry over to this realistic motor task. We now ask which member of this family of task-compatible solutions best matches neural data.

### 2.7 Matching neural data requires an intermediate degree of recurrent restructuring

We have shown that varying *γ* produces a family of networks that all generate accurate muscle outputs but differ in their internal dynamics. We now ask whether any of these networks resemble the neural circuits that actually produce reaching movements.

We compared RNN population-level activity to simultaneously recorded M1/PMd neural data in a ±400 ms window around movement onset. We used centered kernel alignment (CKA) to measure similarity between the population-level correlation structure of the RNN and that of the neural recordings (Fig. 5A) [28, 56]. CKA is a natural metric in the context of our theory, since the correlation function is precisely the population-level quantity that the DMFT determines. Singular vector canonical correlation analysis, the metric used by Sussillo et al. [27], yields a similar picture, as does omitting the *z*-scoring of RNN and M1/PMd activities applied in Fig. 5A (Sec. SI.6, Fig. SI.5).

**Figure 5:**
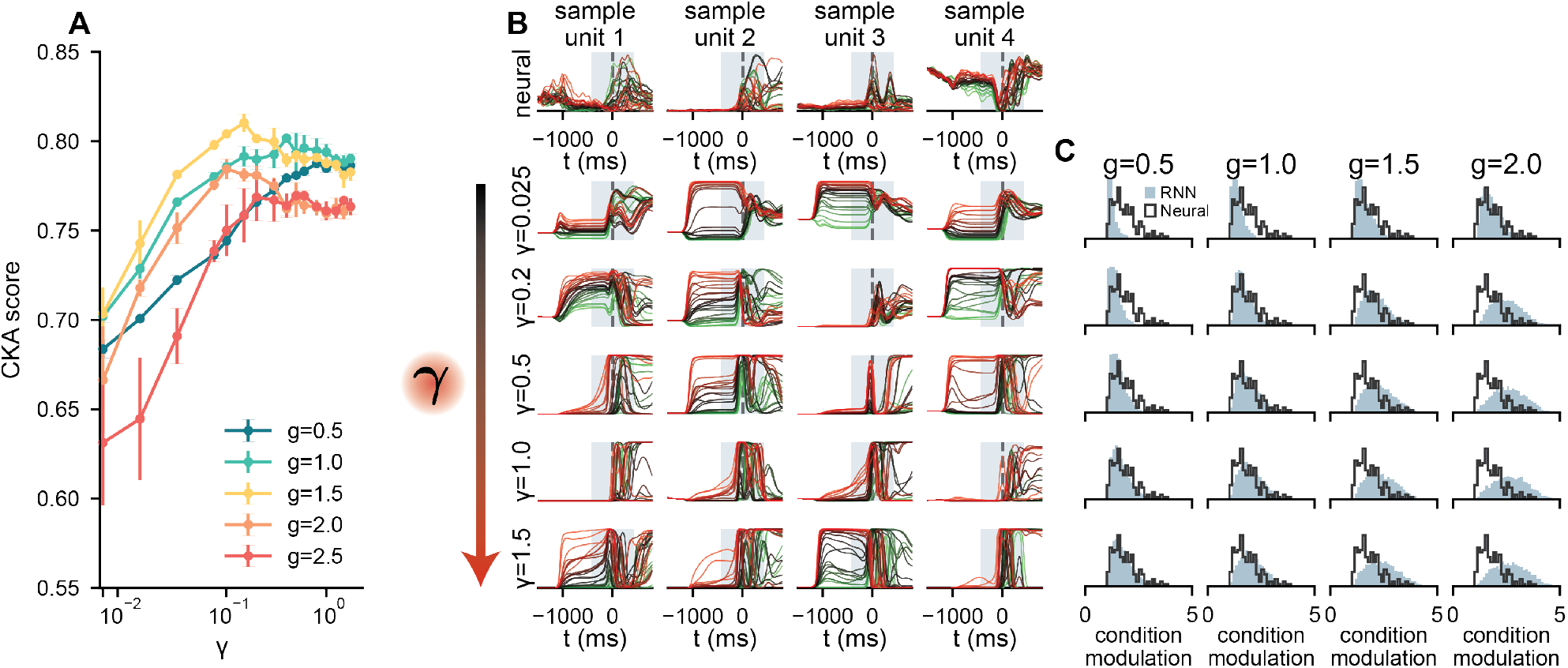
Matching neural data requires an intermediate degree of recurrent restructuring. **(A)** CKA score between RNN and M1/PMd population-level activity as a function of *γ* for different values of *g* (colors), computed in a ± 400 ms window around movement onset. Error bars show standard deviation across runs. **(B)** Example single-neuron responses from M1/PMd recordings (top row) and RNN neurons at *g* = 1 for various values of *γ* (rows below), with four sample neurons per row. Traces show all 27 reach conditions, colored green to red by preparatory activity. The shaded band indicates the ± 400 ms movement window. **(C)** Distributions of per-neuron condition modulation (participation ratio of each neuron’s condition-by-time response matrix) for the same *γ* values as in (B) and different values of *g* (columns), comparing RNN (filled histograms, pooled across 5 runs) and M1/PMd recordings (black outlines).

Similarity to neural data increases sharply with *γ* at small values, confirming that reservoir networks poorly match the structure of motor-cortex responses. For most parameter configurations with *g >* 1, the curves peak at *γ* of order 0.1 and decline at larger values. This nonmonotonicity is more pronounced for some similarity metrics than others (Sec. SI.6). The robust finding is that an intermediate degree of recurrent restructuring substantially improves the match to neural data relative to the reservoir limit, while the largest values of *γ* do not improve it further and can worsen it.

We next examined the single-neuron correlates of this population-level alignment. Our theory predicts that individual neurons are i.i.d. samples from a distribution over response profiles whose character is controlled by *γ*, and that recurrent restructuring drives this distribution away from Gaussianity in a task-dependent manner. Fig. 5B displays example single-neuron responses from M1/PMd recordings alongside those from RNNs at several values of *γ* for *g* = 1. At small *γ*, RNN neurons show little differentiation across reach conditions, with similar temporal profiles regardless of which reach is performed, reflecting the weak condition-based modulation provided by a marginally stable reservoir. As *γ* increases, neurons develop richer condition-dependent structure, including differentiated ramping activity during the delay period and diverse temporal profiles across conditions, increasingly resembling the responses of the M1/PMd neurons.

We quantified this by computing, for each neuron, the participation ratio of the singular values of its condition-by-time response matrix (this per-neuron measure is distinct from the population PR defined above). High values indicate condition-dependent modulation; low values indicate approximately condition-invariant activity. The distributions of this quantity across both M1/PMd and RNN neurons are shown in Fig. 5C. For RNNs, at small *γ*, and particularly for *g* ≤ 1, these distributions are concentrated at low values, reflecting weak condition modulation. As *γ* increases, the distributions broaden and shift rightward, more closely resembling the distribution observed in M1/PMd recordings. At the largest values of *γ*, the distributions overshoot the neural reference for *g >* 1 more substantially than they undershot it at small *γ*, paralleling the decline in population-level similarity seen for *g >* 1 in Fig. 5A. The same intermediate regime of recurrent restructuring that best matches neural activity at the population level also best reproduces the statistics of individual neurons.

## 3 Discussion

We have developed a theoretical framework for task-trained RNNs in which a single parameter, *γ*, controls the degree to which learning reshapes recurrent connectivity. Our DMFT characterizes both the population-level and single-neuron consequences of this restructuring, showing that population-level quantities converge to a deterministic limit while individual neurons are i.i.d. samples from a (generally non-Gaussian) distribution whose character reflects the balance of randomness and structure in the underlying connectivity. Applied to simplified tasks, the theory yields analytical and numerical predictions for how recurrent restructuring amplifies task-relevant frequencies, suppresses chaos, and enables temporal generalization. Applied to a neuroscientifically motivated motor task, the framework generates a continuous family of networks that all produce accurate muscle outputs but differ in their internal dynamics, and we find that matching motor-cortex recordings requires an intermediate degree of recurrent restructuring.

These results bear on a basic question about neural circuits, namely how much learned structure they contain relative to disorder. The pervasive heterogeneity of single-neuron responses across many brain areas is consistent with a large degree of randomness in connectivity, yet the structured, low-dimensional population-level dynamics observed in motor cortex and elsewhere require some task-relevant recurrent structure. Our motor-cortex comparison is consistent with a picture in which this circuit is largely random but contains a small degree of learned recurrent structure sufficient to support the required dynamics. Whether this picture extends to circuits underlying cognitive tasks, where compositional and generalizable representations are essential [21, 22], is an important open question that our framework is well-positioned to address.

A recurring challenge when using task-trained RNNs as models of neural circuits is that each trained network yields a single internal solution, and it is unclear whether its resemblance to neural data reflects the structure of the task or incidental features of the optimization. To make progress, one needs a method for generating multiple task-compatible solutions that satisfies two desiderata. First, the source of diversity should be a controllable parameter with reproducible effects. Second, that parameter should have an interpretable impact on the learned representations, so that when a particular solution matches neural data, one can draw specific conclusions about what properties of the network were responsible.

Existing approaches fall short on one or both counts. Ad hoc regularization, of the type used in [27], provides some control over the learned solution, but without a theory linking regularization to its effects on internal dynamics, interpretation remains difficult [28]. Qian and Pehlevan [57] proposed a more systematic method in which previously found solutions are iteratively projected out to encourage the discovery of qualitatively distinct networks. This explores the solution space more thoroughly, but at considerable computational cost, and the structure of each successive solution is defined only by its dissimilarity from prior ones rather than by an interpretable parameter. Perhaps the most common approach is to train small networks with different random seeds, exploiting finite-size fluctuations to produce diverse solutions [28, 58–60]. However, such finite-size diversity is neither controllable nor interpretable.

Our framework satisfies both desiderata. The parameter *γ* continuously and controllably varies the degree of recurrent restructuring, and the DMFT provides an explicit theory of how it reshapes temporal correlations and single-neuron response distributions. When a particular value of *γ* best matches neural data, this directly quantifies the degree to which task learning has restructured recurrent connectivity in the circuit being modeled (as is evident, for example, when examining the eigenvalues of the learned connectivity; Fig. 4D,E). Importantly, this family of solutions persists in the large-*N* limit, providing a mechanism for diversity that does not rely on finite-size effects. We speculate that finite-size effects are unlikely to be a meaningful generator of diversity in real mammalian circuits. The human cortex contains roughly 10^10^ neurons, and while the more relevant quantity for scaling is the number of inputs per neuron (roughly 10^4^, which equals *N* in our fully connected networks), this is still large. If finite-size fluctuations were the relevant source of variability, brains this large should self-average toward a single prototypical solution. That individuals differ enormously in their behavioral strategies and cognitive styles suggests more structured mechanisms are at work, of which differences in the degree of recurrent restructuring provide one concrete example.

In related work, several papers considered feedforward networks in the lazy and rich regimes to understand the learned neural representations and algorithms underlying phenomena like context-dependent decision making [61], multitask cognition [62], equality reasoning [63], and the influence of the rank of the initial connectivity on learned representations [64]. For RNNs, Huang et al. [60] empirically studied solution degeneracy by varying network size, task complexity, regularization, and feature-learning strength (also using a *µ*P parameterization with parameter *γ*), and mapped how each factor shapes the space of solutions. Our theory focuses on the high rank regime and provides a precise account of what converges and what does not in the large-*N* limit. Population-level quantities such as the correlation function become deterministic, but a large degenerate manifold of weight-space solutions remains, across which individual weights and single-neuron responses vary. Exploring the structure of this manifold, as done by Huang et al. [60], is an interesting avenue for future work; we return to a related point when discussing representational drift below.

Schuessler et al. [65] observed that varying the magnitude of the readout weights, which corresponds to varying *γ*, leads to a transition between “lazy” and “rich” learning regimes, that is, a reservoir regime and a regime with recurrent restructuring. Focusing on the extremes, they identified the rich regime with an aligned readout in which the readout weights lie in the subspace spanned by the top principal components of activity, and the lazy regime with an oblique readout where they do not. They provided a mean-field account of these regimes for noise-driven linear networks in a stationary state, building on earlier work considering tasks depending on the fixed point of a linear RNN [66]. Some of the present authors subsequently analyzed the gradient flow learning dynamics of linear, non-stationary, noise-driven RNNs [67], focusing on the contrast between outlier eigenvalue emergence in an ultra-rich regime and vanishing dynamical change in a very lazy regime. In addition to being limited to linear systems, none of these works showed how solutions vary at intermediate values of *γ*, nor did they consider general sequence-to-sequence tasks. Our results reveal a wide array of behaviors in the (*g, γ*) plane, where larger *g* enhances the chaotic activity of the reservoir while *γ* counteracts it by introducing learned structure.

Our framework connects to, but differs from, the well-studied class of low-rank RNN models [68]. In our energy *E*(**Θ**), the Frobenius norm 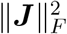 is a factor of *N* larger than all other terms, so that in the learned system the entries *J*_*ij*_ are only an 𝒪(1*/N*) perturbation away from being i.i.d. Gaussian with variance *g*^2^*/N*. Such a small perturbation might seem negligible, but if it has coherent structure, it can produce 𝒪(1) macroscopic effects, pulling outlier eigenvalues out of the random bulk and reshaping activations and dynamics. This is precisely what we observe, suggesting that the learned weight matrix resembles an i.i.d. Gaussian matrix plus a low-rank correction encoding task-relevant structure. In the random-plus-low-rank connectivity literature, this decomposition is specified by construction, either with the low-rank part independent of the random bulk [68, 69] or correlated with it [70, 71]. In our setting, the decomposition is never specified but instead emerges implicitly from end-to-end training; at *β* → ∞, the learned weights can be understood geometrically as uniform samples from the intersection of the high-dimensional spherical shell corresponding to i.i.d. Gaussian weights with a low-dimensional, nonlinearly curved manifold defined by the constraint that the network solves the task. This difference in how the weights are determined leads to a corresponding difference in the DMFT. In random-pluslow-rank models, the order parameters consist of two-time correlation functions tracking bulk-induced fluctuations together with a finite number of scalar overlaps with the low-rank directions [72]. In our theory, the same bulk correlation functions appear, but the role of the low-rank overlaps is replaced by the non-Gaussian tilting of the single-site measure, which implicitly encodes whatever low-rank structure the task demands. Making this correspondence precise is an interesting direction for future work.

During preparation of this manuscript, Bauer et al. [73] released a preprint on feature learning in RNNs, using a mathematical framework largely shared with both our earlier conference presentation [37] and the present work. Despite this common formalism, their work and ours address different questions and operate in different regimes. Their central question is whether and when weight sharing across timesteps in an RNN matters relative to a deep feedforward network with untied weights, a comparison studied primarily through endpoint-supervised tasks in which input enters at the first timestep and a prediction is read out at the last. No such comparison arises in our setting, where the task depends on RNN activity at all time points, as is typical when training RNNs. Their analytical results are restricted to linear activations, and their numerical validation of the nonlinear theory is limited to a regime where learned representations differ only mildly from the reservoir baseline; linear networks cannot exhibit chaos or generate autonomous dynamics. By contrast, we solve the full nonlinear DMFT and demonstrate phenomena inaccessible in the linear or near-reservoir regime, including chaos suppression, the phase transition to low-dimensional dynamics, and temporal generalization. Finally, no analog of our parameter *γ* appears in their framework; the degree of feature learning is an implicit consequence of task signal strength rather than a continuously controllable quantity, precluding the kind of systematic exploration of the solution space and comparison to neural data that *γ* enables here. The two works are thus complementary in focus.

Our finding that networks matching neural data require non-Gaussian single-neuron statistics connects to recent work on statistical models of neural populations. Gaussian models of neural activity, or simple nonlinear transformations thereof, have been surprisingly successful in capturing the statistics of systems such as head-direction cells [7] and hippocampal place fields [8]. In the present setting, however, the reservoir regime is Gaussian and fails to match neural data. Recurrent restructuring provides a systematic, theoretically grounded way to go beyond Gaussian models, with *γ* continuously controlling the degree of non-Gaussianity. Comparing the single-neuron distributions produced by recurrent restructuring with those measured in systems where firing rates or membrane potentials are known to be strongly non-Gaussian [74, 75] could be a productive direction for future work.

Our theory also makes predictions about variability over the timescale of learning. Once the Langevin dynamics reach equilibrium, the weights continue to explore configurations sampled from the Gibbs distribution. Individual weights and single-neuron responses evolve continuously over the learning time *s*, but population-level quantities such as the correlation function and network outputs remain stable, up to fluctuations that shrink as 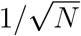. For large networks undergoing noisy plasticity, behavior would thus be stable despite continual turnover of single-neuron activity [55, 76]. This is reminiscent of representational drift, the empirical phenomenon in which the response properties of single neurons change continuously over days and weeks while network-level task performance remains stable [77, 78]. Past work has hypothesized that recurrent dynamics play an important role in determining the relative stability of representations of different stimuli [78]. Characterizing the equilibrium and non-equilibrium dynamics over the learning timescale in our framework, as has been done for feedforward networks [79, 80], could allow one to isolate the role of recurrent dynamics in a theoretically precise way.

Our framework opens several directions for further investigation. The compositional and modular representations that emerge when RNNs are trained simultaneously on multiple cognitive tasks [21, 22, 81] clearly require recurrent restructuring, and studying how *γ* controls the emergence of such structure is a natural next step. Applying the theory across a wider range of motor, decision-making, and cognitive tasks would test whether the picture suggested by our motor-cortex results, that circuits are largely random with varying degrees of task-relevant recurrent structure, extends to other systems.

Finally, our results raise the question of why large neural circuits might tend to operate in a regime where recurrent connectivity is only modestly restructured. A possible answer lies in the biological constraints on learning. Training a readout from a fixed representation is straightforward, requiring only a local learning rule that depends on pre- and postsynaptic activity, and corresponds in our framework to learning the readout weights ***V*** while leaving recurrent weights ***J*** unchanged. Restructuring recurrent connectivity, by contrast, requires propagating error signals backward through time, a computation that is difficult for a biological circuit to implement [82], perhaps even more so than the spatial credit assignment problem faced by feedforward networks. If the recurrent credit assignment problem is indeed hard for biology, then neural circuits may prefer solutions that reshape random connectivity as little as possible while still achieving adequate performance. This tendency would be especially pronounced in large mammalian circuits, where the high-dimensional parameter space and the relatively low-dimensional nature of behavioral tasks create a vast degeneracy of solutions, many of which require only modest recurrent restructuring, and selection pressure need only find one.

## Acknowledgements

We are indebted to Mark Churchland for kindly sharing data from Sussillo et al. [27], and for helpful discussions. We thank Juan Carlos Fernández del Castillo for helpful comments on a previous version of this manuscript.

In the initial phase of this work, B.B. was supported by a Google Ph.D. Fellowship, and J.A.Z.-V. and C.P. were supported by NSF Award DMS-2134157, NSF CAREER Award IIS-2239780, and a Sloan Research Fellowship. At present, D.G.C. is supported by the Kempner Institute for the Study of Natural and Artificial Intelligence at Harvard University. B.B. is supported by the Center of Mathematical Sciences and Applications at Harvard University. J.A.Z.-V. is supported by the Office of the Director of the National Institutes of Health under Award Number DP5OD037354, and by a Junior Fellowship from the Harvard Society of Fellows. C.P. is presently supported by NSF CAREER Award IIS-2239780, DARPA grants DIAL-FP-038 and AIQ-HR00112520041, and The William F. Milton Fund from Harvard University. This work has been made possible in part by a gift from the Chan Zuckerberg Initiative Foundation to establish the Kempner Institute for the Study of Natural and Artificial Intelligence.

## Declaration of generative AI and AI-assisted technologies in the manuscript preparation process

During the preparation of this work the authors used Claude Opus 4.6/4.7 and Claude Code in order to write code, prepare figures, and edit and proofread text. After using this service, the authors reviewed and edited the content as needed and take full responsibility for the content of the published article.

## Supplementary Information

### SI.1 Core dynamical mean-field theory

This appendix derives the dynamical mean-field theory (DMFT) for our task-trained RNN. We begin by specifying the order of limits we analyze (Sec. SI.1.1), then carry out the derivation in discrete time for a single input–output sequence (Sec. SI.1.2), and finally describe the numerical methods used to solve the resulting saddle-point equations (Sec. SI.1.3). Subsequent appendices extend the theory to multiple sequences and generalization (Sec. SI.2), and to the continuous-time, time-translation-invariant, and linear regimes (Sec. SI.3).

#### SI.1.1 Which limit do we analyze?

As mentioned in the main text, we consider Langevin dynamics of the RNN parameters **Θ**, with learning time *s* and noise variance 2*/β* (Eq. (4)). We are often interested in the behavior of the system for large *s* (train for a long time), large *β* (inject small amounts of noise), and large *N* (use a large network). However, these three limits generally do not commute [34, 80]. In the present work, we consider the following order of limits.

1. **Large** *s* **limit**. We first take *s* → ∞. In this limit, **Θ** is drawn from the Gibbs distribution ∝ exp(−*βE*(**Θ**)), which depends on *β* and *N*.
2. **Large** *N* **limit**. We next take *N* → ∞. This leads to the DMFT derivation of Sec. SI.1.2.
3. **Large** *β* **limit**. Finally, we can optionally take the low-temperature limit *β* → ∞, in which the task is fit perfectly.

While this order of limits captures the equilibrium behavior, it does not capture the learning dynamics over *s*. We leave an analysis of that regime in the *N* → ∞ limit to future work. It is also distinct from taking the limit *β* → ∞ before the limit *s* → ∞, which would correspond to networks trained using noiseless gradient flow [67].

#### SI.1.2 Derivation of the mean-field equations

We derive the DMFT in discrete time, then take the continuous-time limit in Sec. SI.3.1. We discretize time with step Δ*t*, writing *t*_*n*_ = *n* Δ*t* for *n* = 1, …, *n*_*T*_. The network dynamics read

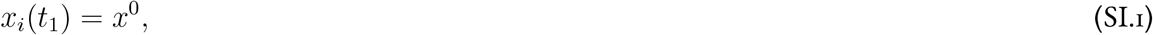

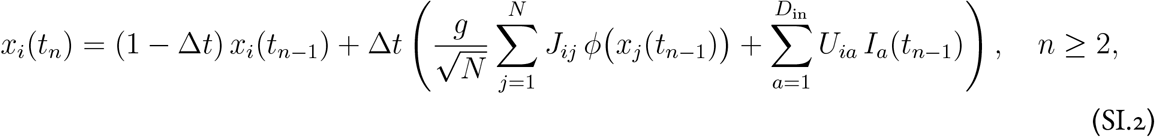

and the readout is

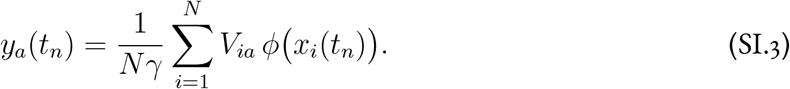

The energy function is

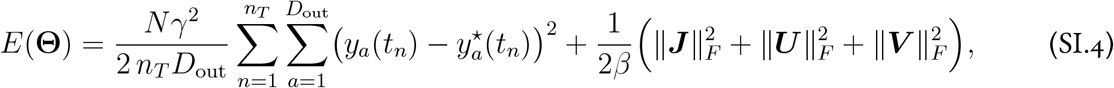

where **Θ** = {***J, U***, ***V***}. Since we consider the *s* → ∞ limit of the Langevin dynamics, we wish to evaluate the partition function for the equilibrium measure of **Θ**,

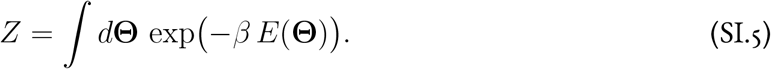

In the energy, the readout *y*_*a*_(*t*_*n*_) depends on **Θ** in a complicated, highly nonlinear way through the network dynamics. To facilitate integration over **Θ**, we use the Martin–Siggia–Rose–De Dominicis– Janssen (MSRDJ) path-integral approach [83–85]. We enforce the network dynamics via delta-function constraints, representing each delta function in its Fourier form with conjugate variables 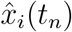 integrated over (−∞, ∞), and similarly introduce auxiliary output fields *ŷ*_*a*_(*t*_*n*_) running over (−*i*∞, *i*∞) for the readout constraint. Averaging over the i.i.d. Gaussian weights **Θ** then gives

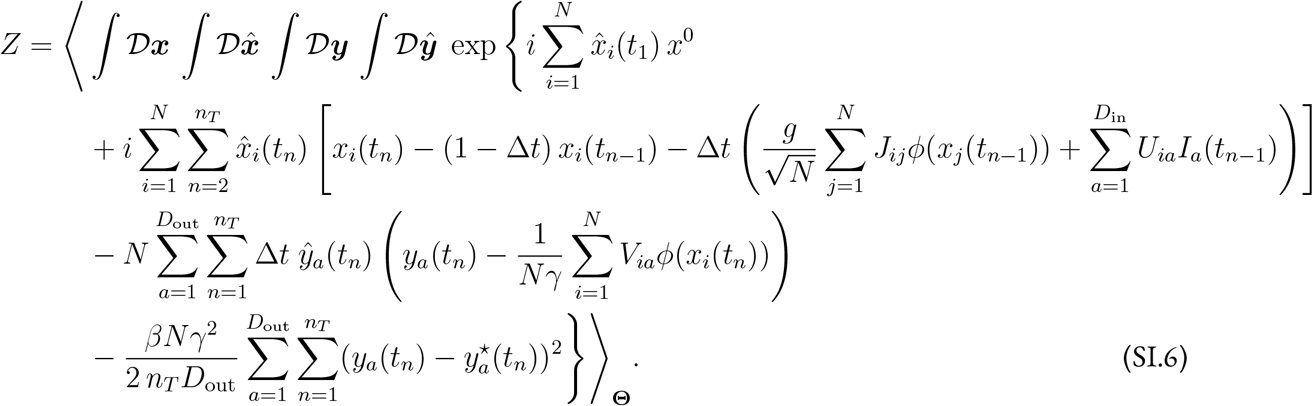

The average over ***J*** generates a term quartic in the activations *ϕ*_*i*_(*t*_*n*_), in which neurons are coupled through the empirical correlation (1*/N*) ∑_*i*_ *ϕ*_*i*_(*t*_*n*_) *ϕ*_*i*_(*t*_*n*_*′*). We decouple this term by promoting the empirical correlation to an order parameter *C*(*t*_*n*_, *t*_*n*_*′*), with conjugate field *Ĉ*(*t*_*n*_, *t*_*n*_*′*) enforcing the definition *C*(*t*_*n*_, *t*_*n*_*′*) = (1*/N*) ∑_*i*_ *ϕ*_*i*_(*t*_*n*_) *ϕ*_*i*_(*t*_*n*_*′*) via a Fourier representation of the delta function. We collect these into *n*_*T*_ × *n*_*T*_ matrices ***C*** (with elements integrated along the real axis) and ***Ĉ*** (along the imaginary axis). The partition function then takes the form

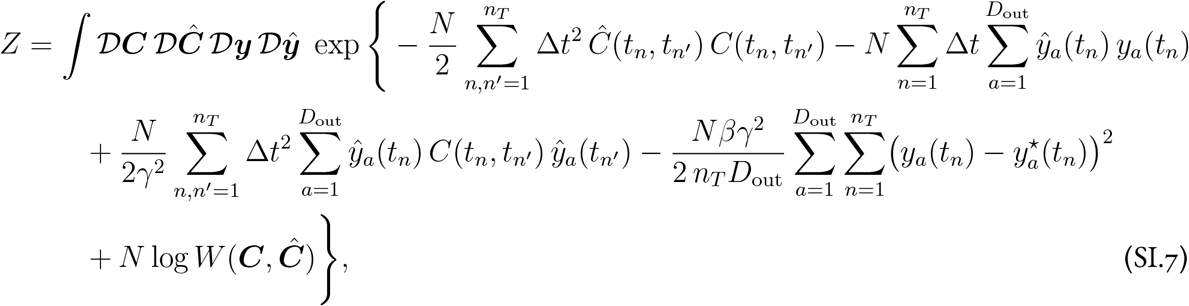

where the single-neuron generating functional *W* (***C, Ĉ***) is defined by

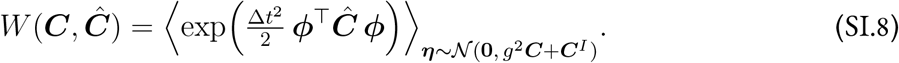

Here 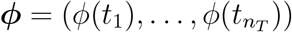 is the *n*_*T*_-dimensional vector of activations generated by the single-neuron dynamics

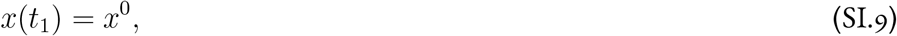

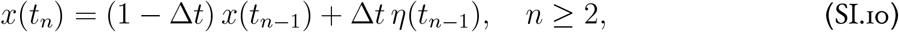

with *ϕ*(*t*_*n*_) = *ϕ*(*x*(*t*_*n*_)), and *η*(*t*_*n*_) is a Gaussian process with covariance ⟨*η*(*t*_*n*_) *η*(*t*_*n*_*′*)⟩ = *g*^2^*C*(*t*_*n*_, *t*_*n*_*′*) + *C*^*I*^(*t*_*n*_, *t*_*n*_*′*). The expectation in Eq. (SI.8) is taken over realizations of *η*(*t*_*n*_).

Reading off the action 𝒮(***C, Ĉ, Y***, ***Ŷ***) such that *Z* = ∫ 𝒟***C*** 𝒟***Ĉ*** 𝒟***Y*** 𝒟***Ŷ*** exp(−*N* 𝒮), we have

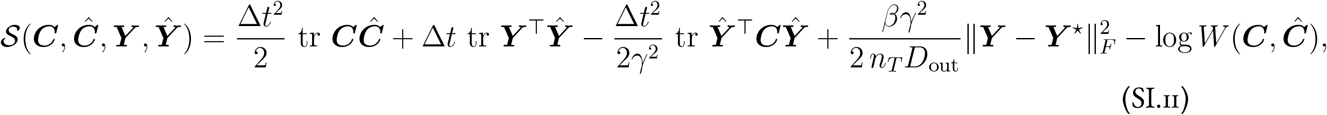

where ***Y*** and ***Ŷ*** are *n*_*T*_ × *D*_out_ matrices with elements *y*_*a*_(*t*_*n*_) and *ŷ*_*a*_(*t*_*n*_) respectively. The action is quadratic in ***Y*** and ***Ŷ***, so their saddle-point evaluations correspond to exact Gaussian integrations.

Evaluating first the saddle point for ***Ŷ*** gives

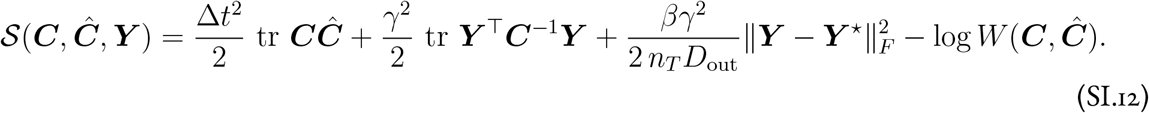

Evaluating next the saddle point for ***Y*** yields the output

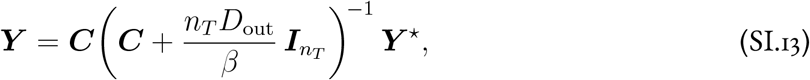

which, in the *β* → ∞ limit, reduces to ***Y*** = ***Y*** ^⋆^, provided that ***C*** is invertible.

Substituting Eq. (SI.13) back and simplifying, the action reduces to

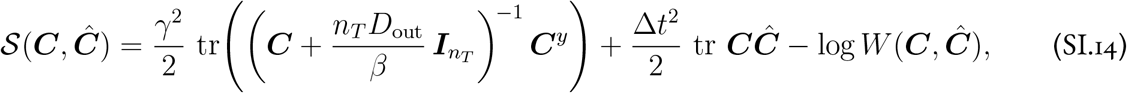

where ***C***^*y*^ is the *n*_*T*_ × *n*_*T*_ target correlation matrix with elements

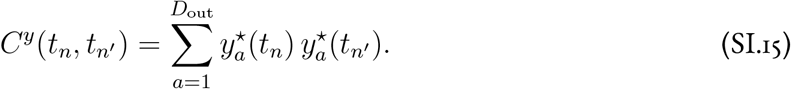

Recall that elements of ***Ĉ*** run over (−*i*∞, *i*∞), and those of ***C*** run over (−∞, ∞). For the saddle-point approximation to be valid, the action must therefore be maximized in ***Ĉ*** and minimized in ***C***. The stationary point of interest is therefore a genuine saddle, with directions of both negative and positive curvature, and solving for this point numerically is nontrivial (Sec. SI.1.3).

We can make the structure of this action more transparent by writing

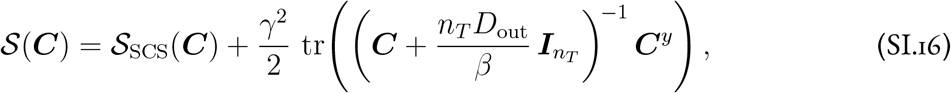

where

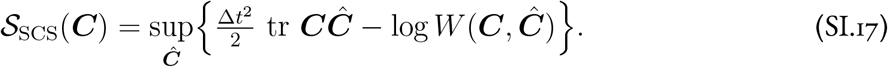

The first term in 𝒮(***C***), 𝒮_SCS_(***C***), encodes the statistics of the random, unstructured network inherited from the SCS theory. The second is the learning term, which penalizes misalignment between the network’s temporal correlations and those of the target.

##### Self-consistency and the tilted measure

The saddle-point condition ∂𝒮*/*∂*Ĉ*(*t*_*n*_, *t*_*n*_*′*) = 0 yields a self-consistency equation that determines *C*(*t*_*n*_, *t*_*n*_*′*) in terms of the single-site measure defined by *W* (***C, Ĉ***). Concretely, *W* (***C, Ĉ***) defines a tilted probability measure on input currents,

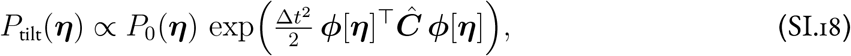

where *P*_0_(***η***) = 𝒩(***η***; **0**, *g*^2^***C*** + ***C***^*I*^) is the untilted Gaussian measure and ***ϕ*** is the activation profile obtained by passing ***η*** through the single-neuron dynamics (SI.9) and applying the nonlinearity (the induced distribution over activations takes the form 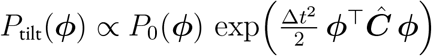, where *P*_0_(***ϕ***) = ∫ *d****η****P*_0_(***η***)*δ*(***ϕ*** − ***η***), noting that the tilting factor is constant where the integrand is nonzero). The self-consistency condition requires that the correlation function computed under the tilted measure reproduces the order parameter,

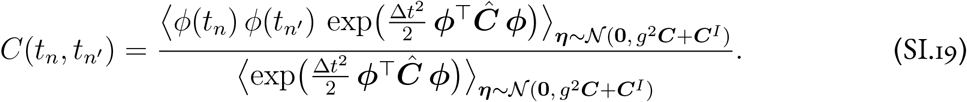

When ***Ĉ*** = **0**, as occurs in the reservoir limit *γ* → 0^+^, the tilting factor reduces to unity, the input current ***η*** is simply Gaussian, and Eq. (SI.19) recovers the classical SCS self-consistency equation. When ***Ĉ*** ≠ 0, the exponential tilting biases the population toward neurons whose input currents produce activations aligned with the structure encoded in ***Ĉ***, driving single-neuron statistics away from Gaussianity in a task-dependent manner. This tilted measure and the associated self-consistency equation will reappear, in progressively generalized forms, in the multi-sequence and generalization settings of Sec. SI.2.

One statistical framework that provides conceptual insight into the adapted single site distribution (and also provides a practical sampling scheme for numerics) is *importance sampling*, where one reinterprets averages ⟨*F* (*η*)⟩_*η*~*p*(*η*)_ over a challenging distribution *p*(*η*) with averages over a simpler distribution *q*(*η*) as 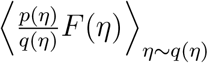 [86]. In our DMFT, we can take *q*(*η*) to be the Gaussian process, while *p*(*η*) is the fully non-Gaussian adapted single site distribution. The *importance weight* 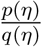 is exactly the tilting factor 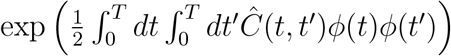.

#### SI.1.3 Numerical solution of the DMFT equations

The DMFT equations must be solved numerically for nonlinear *ϕ*(·). We use an alternating iteration procedure to converge to the saddle point (***C, Ĉ***).

##### Alternating iteration procedure

We assume access to an automatically differentiable implementation [87] of the action 𝒮(***C, Ĉ***). We set hyperparameters 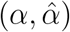 controlling the update speed for ***C*** and ***Ĉ*** respectively, and follow this procedure.

1. Initialize the correlation functions (we use either ***C*** = ***I*** or ***C*** = ***C***_SCS_) and ***Ĉ*** = **0**.
2. Alternate the following updates until a convergence criterion is met:
  - Iterate 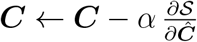 until convergence at fixed ***Ĉ***.
  - Take a single step 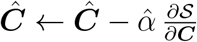.
3. Return the final (***C, Ĉ***).

We find that this scheme is significantly more stable than an algorithm that attempts to exactly compute the Legendre transform sup_***Ĉ***_ 𝒮(***C, Ĉ***) before updating ***C***. To understand why, we next discuss a general perspective on the saddle-point solver in terms of preconditioned gradient flow.

##### A preconditioned flow perspective

To interpret and compare iteration schemes, consider a general preconditioned gradient flow on the DMFT action. Let *ς* represent a flow time and ***P*** a real 2 × 2 preconditioner matrix. The dynamics are

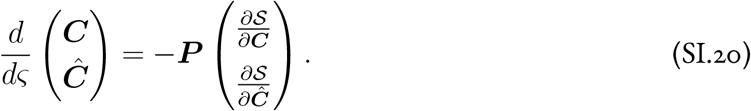

The two natural choices are a diagonal preconditioner ***P***_diag_, which uses ∂𝒮*/*∂***C*** to update ***C*** and ∂𝒮*/*∂***Ĉ*** to update ***Ĉ***; and an off-diagonal preconditioner ***P***_off-diag_, which uses ∂𝒮*/*∂***Ĉ*** to update ***C*** and ∂𝒮*/*∂***C*** to update ***Ĉ***,

**Figure SI.1:**
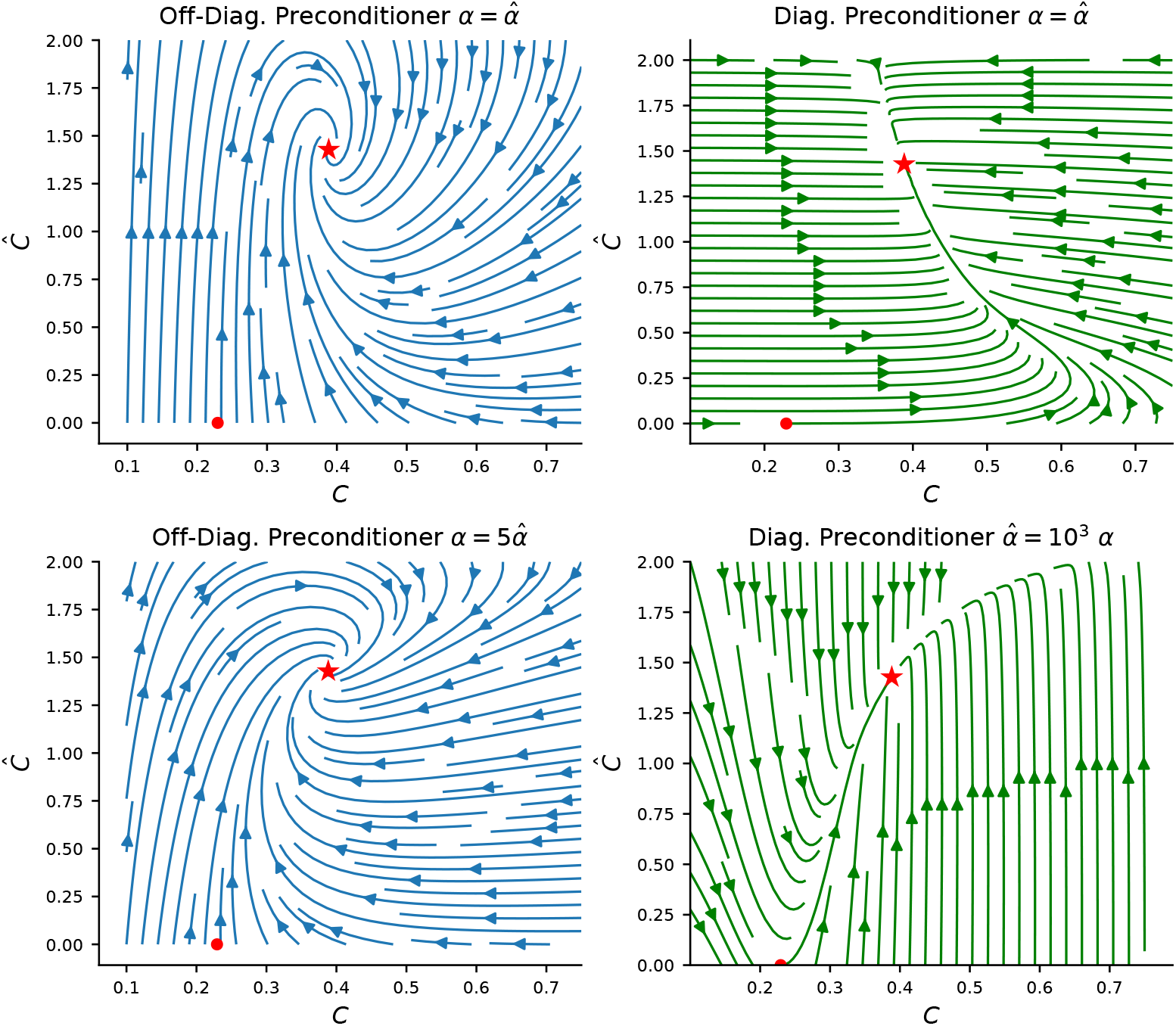
Flow field for preconditioned gradient dynamics for a single frequency *ω* in the time-translation-invariant linear RNN, with (*g, γ, ω*) = (0.9, 0.5, 2). In all cases, the dynamics flow from the SCS solution (red dot) to the DMFT saddle point (red star). The off-diagonal dynamics are better behaved at comparable step sizes 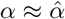, exhibiting mild spiraling near the saddle point. Monotonic convergence in both *C* and *Ĉ* can be achieved in the off-diagonal scheme with 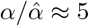, whereas the diagonal scheme requires 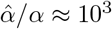.

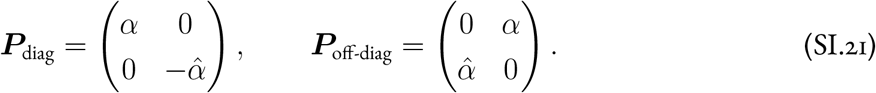

Both choices would in principle converge to the correct saddle point. However, the off-diagonal preconditioner (which is essentially our iterative algorithm) is significantly more numerically stable for non-infinitesimal step sizes. Fig. SI.1 illustrates the convergence dynamics for these two preconditioners in the time-translation-invariant linear RNN setting (Sec. SI.3.3).

##### Computing the differentiable action

We compute 𝒮(***C, Ĉ***) using an importance-sampling scheme that is compatible with automatic differentiation libraries such as PyTorch [88] and JAX [87]. The procedure is as follows.

1. Perform a Cholesky decomposition of ***C*** to obtain a square root ***C***^1*/*2^.
2. Sample *M* random Gaussian vectors 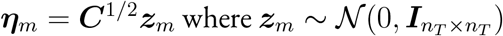.
3. Integrate the difference equation (Eq. (SI.9)) to compute ***x***_*m*_ from ***η***_*m*_.
4. Compute the firing-rate vector ***ϕ***_*m*_ = *ϕ*(***x***_*m*_).
5. Estimate the single-site partition function 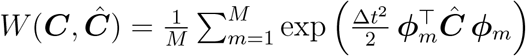.
6. Compute the action 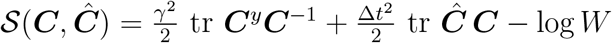.

In the time-translation-invariant regime, the Cholesky decomposition can be replaced by Fourier transforms, reducing computational complexity from 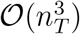 to 𝒪(*n*_*T*_ log *n*_*T*_); the resulting specialized solver for nonlinear networks is described in Sec. SI.3.2.

### SI.2 Multi-sequence settings and generalization

The derivation in Sec. SI.1.2 considered a single input–output sequence. We now extend it to a batch of sequences (Sec. SI.2.1) and to generalization on unsupervised sequences or time intervals (Sec. SI.2.2). Throughout, the structure of the action and self-consistency equations carries over with minimal modification; we use parallel notation to make the correspondence explicit.

#### SI.2.1 Multi-sequence (batch) case

We generalize to a batch of *B* input–output pairs, consisting of input sequences 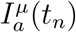, target outputs 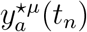, and initial conditions *x*^0*µ*^, where *µ* = 1, …, *B* indexes the sequence.

##### Energy and order parameters

The MSE term in the energy *E*(**Θ**) becomes an average over sequences as well as over time and output channels,

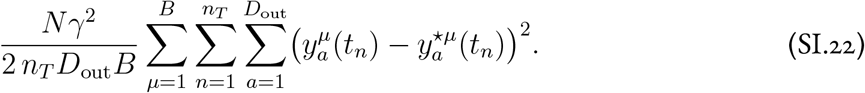

All dynamical variables acquire a sequence index *µ*, and the order parameters are promoted to *C*^*µν*^(*t*_*n*_, *t*_*n*_*′*) and *Ĉ*^µν^ (*t*_*n*_, *t*_*n*_*′*), which carry both temporal and batch indices. To handle these enlarged objects compactly, we introduce Fraktur notation (where ℭ is the Fraktur version of *C*, etc.). We write ℭ and 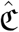 for the full *n*_*T*_ *B* × *n*_*T*_ *B* matrices obtained by treating the pair (*µ, t*_*n*_) as a single composite index. We similarly write 𝔜 and 𝔜^⋆^ for the output and target matrices, and ℭ^*y*^ for the target correlation matrix.

##### Derivation and action

The derivation proceeds exactly as in Sec. SI.1.2, with all traces and inverses now acting on the enlarged *n*_*T*_ *B* × *n*_*T*_ *B* matrices. The learning term 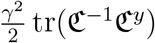 is structurally identical to its single-sequence counterpart, simply promoted to the larger matrices. The only meaningful change appears in the generating functional 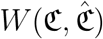, and thus in 𝒮_SCS_(ℭ). Because time does not propagate along the batch dimension, the single-site dynamics consist of *B* independent trajectories, each evolving from its own initial condition *x*^0*µ*^.

The saddle-point equations for 𝔜 and 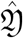 are solved as before. The output is

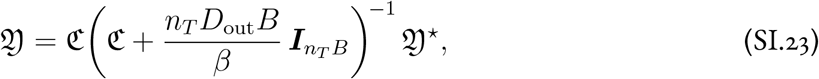

which is the direct analog of Eq. (SI.13), with Fraktur matrices replacing their single-sequence counterparts and the regularization factor reflecting the larger number of supervised data points. After substitution, the action is

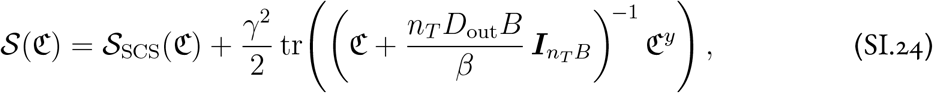

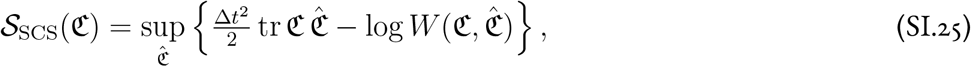

which is the direct analog of Eqs. (SI.16)–(SI.17).

##### Self-consistency

The self-consistency equation likewise generalizes in the expected way. The untilted Gaussian process is now sampled from the full *n*_*T*_ *B* × *n*_*T*_ *B* covariance *g*^2^ℭ + ℭ^*I*^, the tilting is governed by 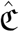, and the self-consistency condition reads

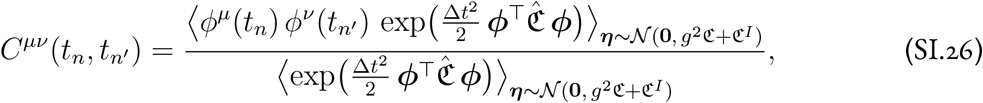

where ***ϕ*** now denotes the concatenation of activations ***ϕ***^*µ*^ across all *B* sequences. This is the direct analog of Eq. (SI.19), with the single-sequence measure replaced by the batched one.

#### SI.2.2 Generalization

The DMFT developed above characterizes the dynamics of the learned network over the sequences at which it was supervised. We now extend the framework to *generalization*, asking what the network dynamics look like on unsupervised (held-out) sequences or time intervals.

##### Setup

We partition the *B* sequences into a training set 𝒯 ⊂ {1, …, *B*} and a test set 𝒯 ^*c*^ = {1, …, *B*}\ 𝒯. The energy *E*^tr^(**Θ**) includes contributions only from training sequences, with the MSE in Eq. (SI.22) averaged over *µ* ∈ 𝒯 rather than over all *B*. We write the corresponding partition function as *Z*^tr^ = ∫ *d***Θ** exp(−*β E*^tr^(**Θ**)). Evaluating *Z*^tr^ via the procedure of the preceding sections yields the training DMFT, which determines the train–train block of the order parameters ℭ^tr^ and 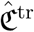.

To compute the dynamical order parameters on the test sequences, we average a separate path integral over the Gibbs measure induced by training,

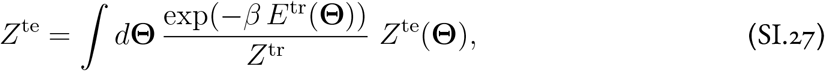

where *Z*^te^(**Θ**) is the path integral that enforces the network dynamics on the test sequences for a given set of weights **Θ**, without any squared-error term. Concretely, *Z*^te^(**Θ**) takes the form

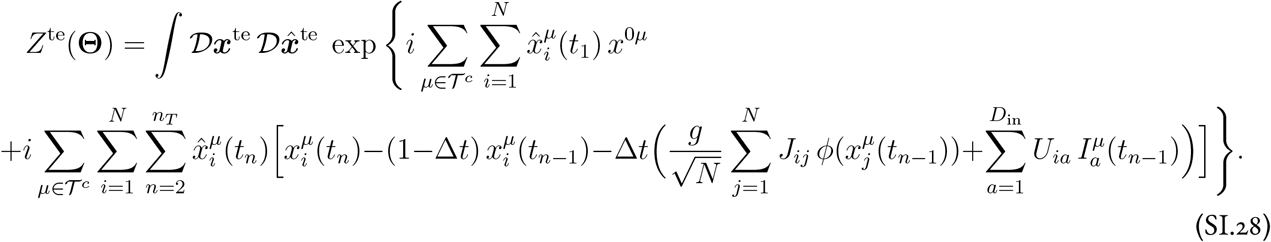

When we average *Z*^te^(**Θ**) over the training Gibbs measure and introduce order parameters for the combined (train, test) system, the Fraktur matrices ℭ, 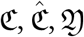, 𝔜, and 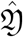, carry indices ranging over all *B* sequences, both train and test. We proceed with the saddle-point calculation as before. The key simplification comes from the structure of the conjugate variables at the saddle point.

##### Vanishing of test conjugate variables

At the saddle point, all components of 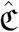 and 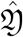 in which at least one index belongs to the test set vanish,

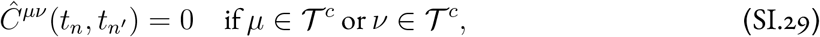

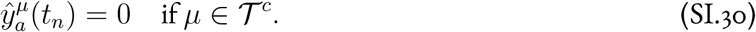

Only the train–train block *Ĉ*^*µν*^ (*t*_*n*_, *t*_*n*_*′*) with *µ, ν* ∈ 𝒯 is nonzero.

This result has a clear physical interpretation. The conjugate variable 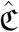 governs the non-Gaussian tilting of the distribution over input currents (Eq. (SI.18)). Test sequences share no error signal, so they cannot tilt one another; the test–test block of 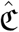 must therefore vanish. Test sequences also play no role in the training objective, so they cannot influence the tilting of training sequences; the train–test and test–train blocks must vanish as well. The only sequences that interact through the tilting mechanism are training sequences, which are coupled by virtue of being used to train the same set of weights **Θ**.

##### Saddle-point solution

As a consequence of Eq. (SI.29), the training order parameters decouple from the test sequences entirely. The train–train block ℭ^tr^ and its conjugate 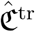 are determined by the same DMFT equations derived in the preceding sections, with no dependence on the test data.

The test-dependent components of ℭ are determined by the same tilted single-site measure, but with the tilting restricted to the train–train block. In the single-site problem, the full (train, test) Gaussian process ***η*** = (***η***^tr^, ***η***^te^) is sampled from the *n*_*T*_ *B* × *n*_*T*_ *B* covariance *g*^2^ℭ + ℭ^*I*^, which couples train and test sequences through their shared dependence on the same weights. Each sequence’s activations ***ϕ***^*µ*^ are obtained by passing the corresponding ***η***^*µ*^ through the single-neuron dynamics (SI.9). However, because only the train–train block of 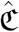 is nonzero, the tilting involves exclusively the training components of ***ϕ***. The self-consistency equation for all components of ℭ takes the unified form

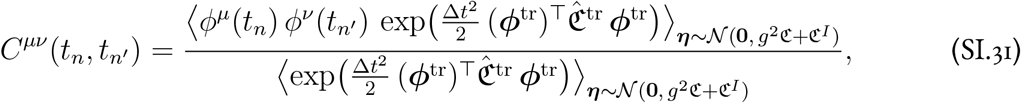

where ***ϕ***^tr^ denotes the concatenation of activations across training sequences and times. Eq. (SI.31) is the direct analog of the batch self-consistency equation (Eq. (SI.26)). The key difference is that the tilting acts only on training activations ***ϕ***^tr^, while the observable *ϕ*^*µ*^(*t*_*n*_) *ϕ*^*ν*^(*t*_*n*_*′*) in the numerator may involve test activations. The equation holds for all pairs (*µ, ν*), whether both are in 𝒯, both in 𝒯 ^*c*^, or one in each.

##### Output

The output formulas likewise parallel those of the preceding sections, with one new element. The train output is given by

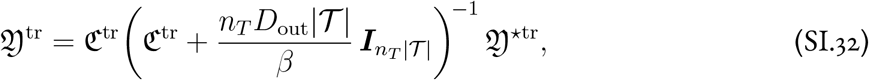

which is the direct analog of Eqs. (SI.13) and (SI.23), restricted to training sequences. The test output takes the form of a kernel regression, with the cross-covariance between test and train sequences mediating the prediction,

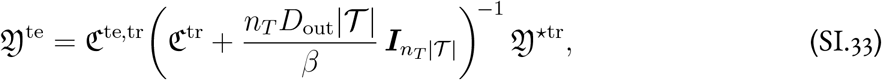

where ℭ^te,tr^ is the *n*_*T*_ |𝒯 ^*c*^| × *n*_*T*_ |𝒯 | cross-covariance block. The only difference from Eq. (SI.32) is that ℭ^tr^ in front of the regularized inverse is replaced by ℭ^te,tr^. The test output is thus determined entirely by the correlation between test and train dynamics and the training targets, with no direct dependence on test targets (which are, by assumption, unavailable).

##### Application to temporal generalization within a single sequence

The same formalism applies not only to held-out sequences but also to held-out time intervals within a single sequence. In the sine wave generation task of Sec. 2.5, for example, the network is supervised over [0, *T*] and we ask what it produces for *t > T*. To handle this, we partition the time axis into a supervised block (*µ* = 1, covering [0, *T*]) and an unsupervised block (*µ* = 2, covering [*T, T*_tot_]), and assign the first to 𝒯 and the second to 𝒯 ^*c*^. The formalism then proceeds exactly as above, with one important difference. Unlike the true batch setting, where each sequence starts from an independent initial condition and time does not propagate across sequences, here time *does* propagate across blocks. The state at the end of the supervised window serves as the initial condition for the unsupervised window, coupling the two blocks in the single-site dynamics. This coupling is what allows the framework to predict whether learned temporal structure (such as oscillations) persists beyond the training interval.

The numerical solver consists simply of the **ℭ** iterations with the tilting matrix fixed. That is, **ℭ**^tr^ is held at its training saddle-point value while the test-dependent components of **ℭ** are iterated.

### SI.3 Continuous time, time-translation invariance, and linear networks

In this appendix we specialize the DMFT to settings that admit simplification and, in some cases, closed-form solutions. We first take the continuous-time limit (Sec. SI.3.1), then consider time-translation-invariant (TTI) solutions that decouple the DMFT across Fourier modes (Sec. SI.3.2), and finally solve the TTI equations analytically for linear networks (Sec. SI.3.3). The appendix concludes with a comparison between linear and nonlinear networks (Sec. SI.3.4), highlighting a qualitatively new phenomenon enabled by the nonlinearity, namely the suppression of unused frequencies.

#### SI.3.1 Continuous-time limit

The continuous-time limit is obtained by taking Δ*t* → 0 with *T* = *n*_*T*_ Δ*t* held fixed. Discrete sums ∑_*n*_ Δ*t* (· · ·) become integrals 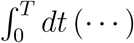. The matrices ***C, Ĉ***, and ***C***^*y*^ become continuous-time kernels *C*(*t,t*′), *Ĉ*(*t,t*′), and *C*^*y*^(*t,t*′). The matrix trace 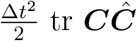 becomes 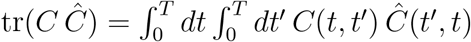, and the learning term 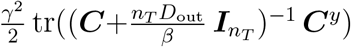 becomes 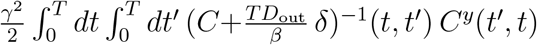, where *δ*(*t* − *t*′) is a Dirac delta. The discrete single-neuron dynamics (SI.9) become (1 + ∂_*t*_) *x*(*t*) = *η*(*t*).

In this limit, the action (SI.16) becomes

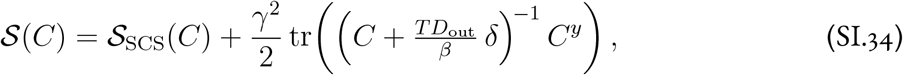

where 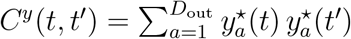 and

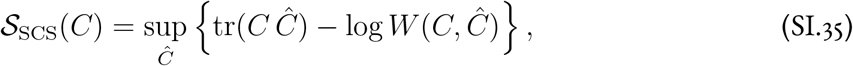

with

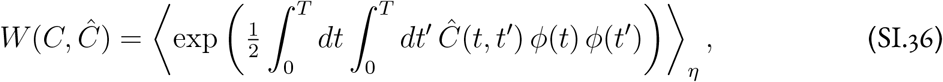

where *ϕ*(*t*) = *ϕ*(*x*(*t*)), *x*(*t*) satisfies (1 + ∂_*t*_) *x*(*t*) = *η*(*t*), and *η*(*t*) is a Gaussian process with covariance *g*^2^*C*(*t, t*′)+*C*^*I*^(*t, t*′). In the *β* → ∞ limit, the regularized inverse reduces to *C*^−1^, the output matches the target exactly (provided *C* is invertible), and the action simplifies to 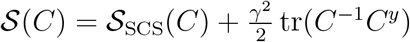. This is the action quoted in Eq. (6) of the main text. The self-consistency equation (Eq. (SI.19)) and the generalizations to the batch (Sec. SI.2.1) and held-out (Sec. SI.2.2) settings carry over with the same substitutions.

#### SI.3.2 Time-translation-invariant solution

A further simplification arises when the system admits a time-translation-invariant (TTI) solution, in which all two-time correlation functions depend only on time differences. Concretely, we assume that the input and target correlations satisfy *C*^*I*^(*t, t*′) = *C*^*I*^(*t* − *t*′) and *C*^*y*^(*t, t*′) = *C*^*y*^(*t* − *t*′), and we seek solutions in which *C*(*t, t*′) = *C*(*t* − *t*′) and *Ĉ*(*t, t*′) = *Ĉ*(*t* − *t*′) likewise. We denote *τ* = *t* − *t*′. (Note that time-translation invariance is a property of the correlation functions, not of the activity *x*(*t*) itself, which will generally fluctuate over time.) This regime is unable to describe inherently nonstationary scenarios such as temporal generalization beyond a finite time window (Sec. 2.5) or a finite-duration motor or cognitive task (Sec. 2.6). The key benefit of this regime is that it enables analysis in the Fourier domain [18], providing both dramatic reductions in computational complexity and a simple spectral interpretation of the learned representations. Thus, this TTI regime is extremely common in DMFT studies [18, 89, 90].

We adopt the Fourier transform convention for Fourier pair *f* (*t*) ↔ *f* (*ω*) as

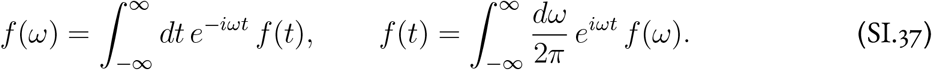

Under TTI structure, the action decomposes in Fourier space as

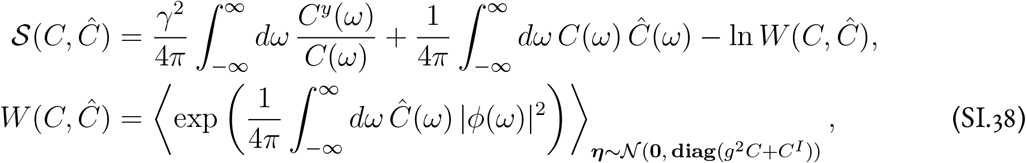

where *ϕ*(*ω*) is the Fourier transform of *ϕ*(*x*(*t*)) with *x*(*t*) generated by the single-neuron dynamics (1 + ∂_*t*_) *x*(*t*) = *η*(*t*). Since *η*(*t*) is a stationary Gaussian process, its Fourier components at distinct frequencies are independent, with *η*(*ω*) having variance *g*^2^*C*(*ω*) + *C*^*I*^(*ω*).

The saddle-point equations ∂𝒮*/*∂*Ĉ*(*ω*) = 0 and ∂𝒮*/*∂*C*(*ω*) = 0 determine *C*(*ω*) and *Ĉ*(*ω*) selfconsistently. The first gives a self-consistency equation for *C*(*ω*) as an expectation under the tilted measure, paralleling Eq. (SI.19),

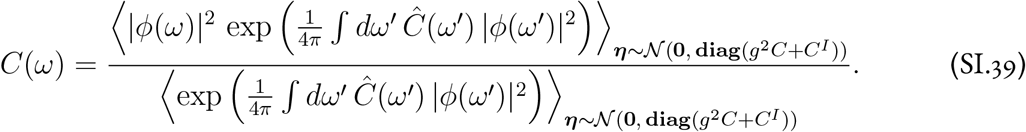

The second gives *Ĉ*(*ω*) in terms of *C*(*ω*) and a functional derivative of log *W* (*C, Ĉ*),

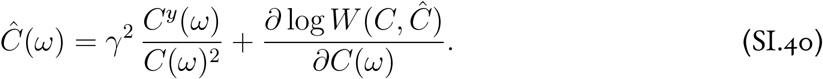

Whether these equations couple different frequencies or decouple into independent problems at each *ω* depends on whether the generating functional *W* (*C, Ĉ*) factorizes across frequencies. The tilting factor in *W* (*C, Ĉ*) is 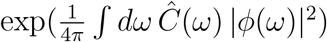, and the expectation is over a Gaussian process with Fourier components *η*(*ω*), which are independent across frequencies. If *ϕ*(*ω*) were a function only of *η*(*ω*) at the same frequency, then |*ϕ*(*ω*)|^*2*^ would be independent across frequencies under the *η*-distribution, the exponent would be a sum of independent terms, and *W* (*C, Ĉ*) would factorize into a product of single-frequency contributions. Both saddle-point equations would then decouple across *ω*.

This is exactly what happens in a linear network, where *ϕ*(*ω*) = *x*(*ω*) = *η*(*ω*)*/*(1 + *iω*), and we exploit it to obtain closed-form solutions that decouple over frequencies *ω* in Sec. SI.3.3. When *ϕ*(·) is nonlinear, however, *ϕ*(*t*) = *ϕ*(*x*(*t*)) is computed pointwise in the time domain, and Fourier-transforming back gives

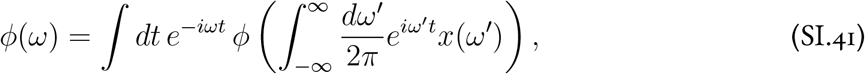

which is a complicated nonlinear mixture of *x*(*ω*) at different frequencies. Thus, *W* (*C, Ĉ*) does not factorize, and both saddle-point equations couple all frequencies together. This inter-frequency coupling is what enables the suppression of task-irrelevant frequencies discussed in Sec. SI.3.4.

##### Spectral bias of the learning term

The structure of Eq. (SI.40) is the frequency-domain counterpart of the general (*t, t*′) principle discussed in Sec. 2.2 of the main text, where the learning term 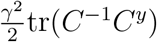 penalizes misalignment between the network’s temporal correlations and those of the target. In the TTI setting, this principle becomes particularly transparent. The ratio *C*^*y*^(*ω*)*/C*(*ω*)^2^ measures how much each frequency is under-represented in the network activity relative to the target, and appears in the equation for *Ĉ*(*ω*) with a prefactor *γ*^2^. When *C*(*ω*) already has substantial power at frequency *ω*, the contribution to *Ĉ*(*ω*) is small, and the tilting of the single-neuron distribution at that frequency is correspondingly weak. When *C*(*ω*) is small relative to *C*^*y*^(*ω*), the contribution to *Ĉ*(*ω*) is large, driving substantial restructuring.

This has two immediate consequences. First, networks trained on higher-frequency targets will generally exhibit larger departures from the random-network correlation structure, because the power spectrum of a random network decays at high frequencies, making the mismatch *C*^*y*^(*ω*)*/C*(*ω*)^2^ large there. Second, networks with larger gain *g* will (at fixed *γ*) show smaller departures, because a larger *g* slows the spectral decay of *C*(*ω*) and reduces the mismatch. Both consequences apply regardless of whether the nonlinearity *ϕ*(·) is the identity (as in the linear theory below) or a saturating function such as tanh. The linear and nonlinear cases differ, however, in whether the saddle-point equations decouple across frequencies; we return to this distinction in Sec. SI.3.4.

##### TTI numerical solver

To solve the TTI saddle-point equations numerically, we use the alternating iteration procedure described in Sec. SI.1.3, maintaining a one-dimensional grid of values for *C*(*ω*) and *Ĉ*(*ω*). The action (SI.38) and its gradients are estimated via importance sampling. Rather than storing and manipulating the full *n*_*T*_ × *n*_*T*_ matrices ***C*** and ***Ĉ***, we sample complex Gaussians *η*(*ω*) with variance *g*^2^*C*(*ω*) + *C*^*I*^(*ω*) at each frequency, inverse-Fourier-transform to obtain *η*(*t*), integrate the single-neuron dynamics to produce *x*(*t*), apply the nonlinearity to get *ϕ*(*t*), and Fourier-transform back to obtain *ϕ*(*ω*). The tilting factor 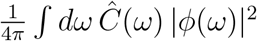 is then computed directly in the frequency domain. This reduces computational complexity from 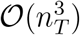 (for a Cholesky decomposition) to 𝒪(*n*_*T*_ log *n*_*T*_) (for Fourier transforms).

#### SI.3.3 Linear time-translation-invariant solution

For a linear network with *ϕ*(*x*) = *x*, the TTI action simplifies considerably. The single-neuron dynamics (1 + ∂_*t*_) *x*(*t*) = *η*(*t*) are a linear filter with transfer function *R*(*ω*) = 1*/*(1 + *iω*), so *x*(*ω*) = *R*(*ω*) *η*(*ω*) and *ϕ*(*ω*) = *x*(*ω*). The generating functional *W* (*C, Ĉ*) can then be computed in closed form, reducing the action to an integral over frequencies of a primitive action at each *ω*,

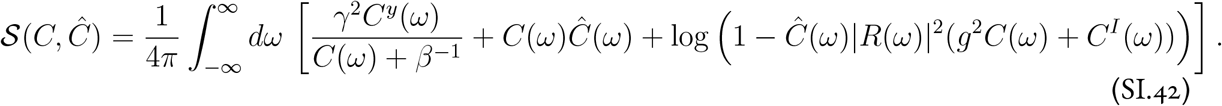

A crucial simplification relative to the nonlinear case is that this action decouples across frequencies, with the saddle-point equations at each *ω* forming a closed system independent of other frequencies. This is because the linear activation *ϕ*(*x*) = *x* does not mix Fourier modes.

Varying with respect to *C*(*ω*) and *Ĉ*(*ω*), we obtain

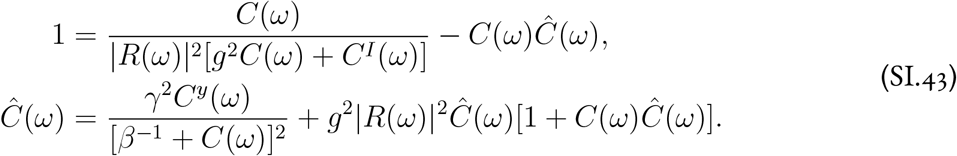

In the reservoir limit *γ* → 0^+^, we have *Ĉ*(*ω*) = 0, and *C*(*ω*) reduces to

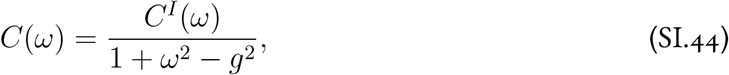

using |*R*(*ω*)|^*2*^ = 1*/*(1 + *ω*^2^). This is the expected result for a random linear reservoir [18, 67].

##### Analytical solution at *γ >* 0

We now solve the coupled equations for *γ >* 0 in the *β* → ∞ limit. Defining

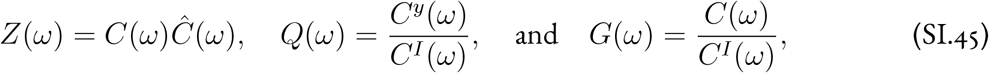

we can rewrite the equations as

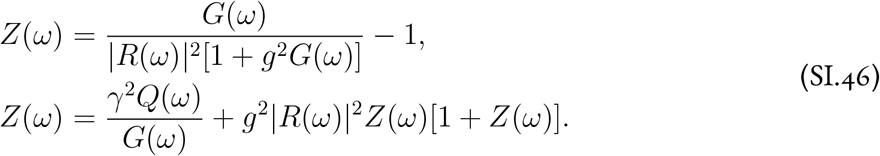

Substituting the first equation into the second and simplifying, we obtain the quadratic equation

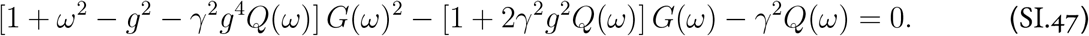

This leads to

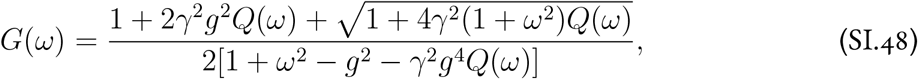

which, upon substituting the expressions for *G*(*ω*) and *Q*(*ω*), gives the closed-form solution quoted in the main text (Eq. (7)). This solution is the physical one because it satisfies 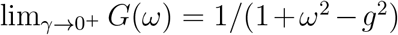, matching the random-network limit; the solution with the negated square root yields 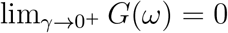 and is unphysical.

Finally, we can solve for

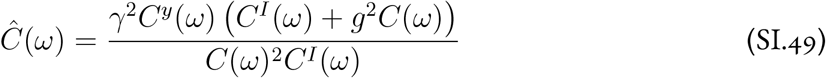

which is manifestly non-negative for any *γ*.

At a conceptual level, this square-root solution closely resembles the solution for the hidden-layer feature kernel of a single-hidden-layer linear Bayesian neural network [44]. In this dynamical setting, however, the assumption of statistical stationarity means that the input and target autocorrelation functions are co-diagonalizable (by the Fourier transform), which is not guaranteed in the feedforward setting, where one must solve a matrix quadratic equation.

##### Stability condition

The solution (SI.48) is free of poles only if

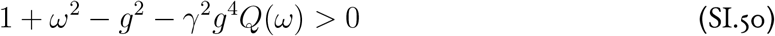

for all *ω*, which requires

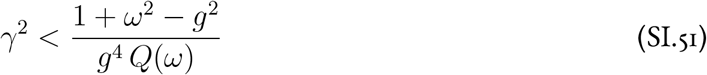

for all *ω*.

##### Interpretation

The ratio *Q*(*ω*) = *C*^*y*^(*ω*)*/C*^*I*^(*ω*) measures the over-representation of a given frequency in the target signal relative to the input. With nonzero *γ*, the transfer function is reshaped to amplify precisely those frequencies. This is made manifest by the small-*γ* expansion,

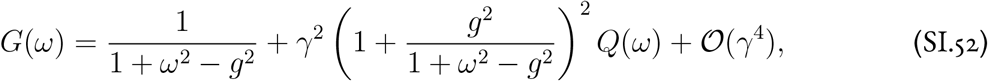

which shows that to leading order in *γ*, each frequency is amplified in proportion to *Q*(*ω*).

As a further illustration, consider white-noise input (*C*^*I*^(*ω*) = 1) and an Ornstein–Uhlenbeck target (*C*^*y*^(*ω*) = 1*/*(1 + *ω*^2^)). Then the solution simplifies to

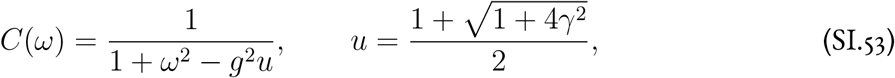

so that feature learning has the effect of increasing the effective gain of the linear reservoir from *g*^2^ to *g*^2^*u*. Stability requires *g*^2^*u <* 1, or equivalently *γ*^2^ *<* (1 − *g*^2^)*/g*^4^.

##### Participation ratio

One useful summary statistic that can be computed from the power spectrum is the effective single-neuron inverse timescale, defined in the stationary setting by (see also Sec. SI.4)

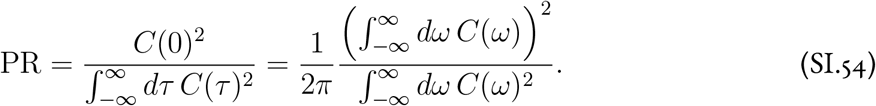

For an Ornstein–Uhlenbeck process with correlation time *τ*,

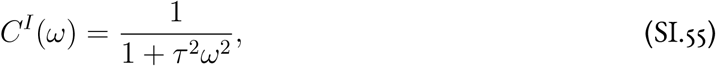

which has participation ratio 1*/τ*. This is the basic reason why we refer to this measure as an inverse timescale, or rate.

If we drive a linear reservoir network with white noise, the single neurons obey Ornstein–Uhlenbeck statistics, and we have

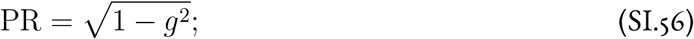

the effective timescale thus diverges as *g* → 1^−^. This is consistent with the result that the participation ratio of activity in linear RNNs should vanish as *g* → 1^−^ in Hu and Sompolinsky [46], and also with the results of Bordelon et al. [67].

Closed-form evaluation of the integrals defining the effective rate for a rich network is not so simple. For the sake of tractability, we first consider a case in which the input process is white noise (*C*^*I*^(*ω*) = 1), such that the two integrals of interest are

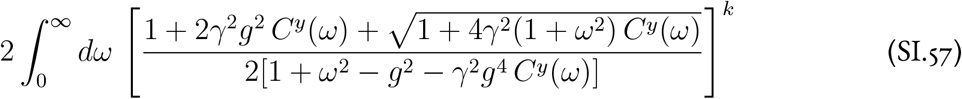

where *k* ∈ {1, 2}. What remains is to make an appropriate choice of *C*^*y*^(*ω*) so that the result is interesting while the computation remains tractable. An interesting test case is the Cauchy (or Lorentzian) spectrum

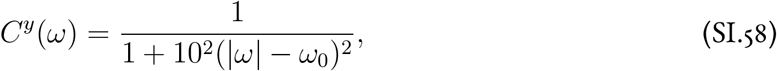

because one can distinguish the peak at *ω*_0_ from the flat background spectrum of *C*^*I*^(*ω*) = 1. However, the resulting integral in general has the form of an elliptic integral.

We therefore resort to numerical evaluation. To do so stably, we make use of the fact that the integrand can be simplified algebraically to

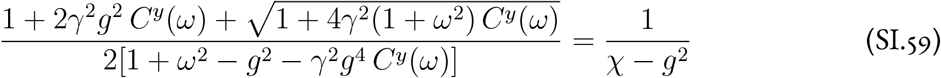

where

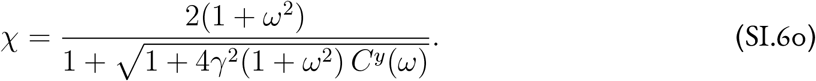

This form is stable numerically even when *C*^*y*^(*ω*) is small. The stability condition is that

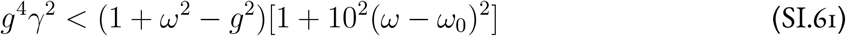

for all *ω*, which is easy to verify numerically. To generate the plots in Fig. 2, we use the adaptive quadrature method provided by `mpmath.quad` to compute the frequency-space integrals.

#### SI.3.4 Frequency suppression in nonlinear networks

In the linear TTI theory derived above, the saddle-point equations decouple across frequencies, with each *C*(*ω*) and *Ĉ*(*ω*) determined independently by the input and target power spectra at that same frequency *ω*. This is because the linear activation *ϕ*(*x*) = *x* does not mix Fourier modes. As a consequence, *Ĉ*(*ω*) ≥ 0 at all frequencies (Eq. (SI.49)), so that learning can only amplify task-relevant frequencies, which is sufficient because the linear activation does not spread power to unwanted frequencies.

The nonlinearity *ϕ*(·) = tanh(·), by contrast, mixes frequencies together. The pointwise application of *ϕ*(·) in the time domain globally reshapes the power spectrum, producing continuous spectral content with peaks at odd harmonics of the fundamental (as can be seen in Fig. SI.2). As a result, the saddlepoint equations for the nonlinear network do not decouple across *ω*, and the solution at one frequency depends on the solution at all others. This inter-frequency coupling enables a phenomenon with no linear counterpart, namely the suppression of frequencies not required by the task. In the nonlinear DMFT solutions (Fig. SI.2), the conjugate order parameter *Ĉ*(*ω*) is positive near the target frequency *ω*_⋆_, but broadly *negative* elsewhere, suppressing task-irrelevant spectral content. Within this negative region, the odd harmonics 3*ω*_⋆_, 5*ω*_⋆_, … appear as local peaks, less suppressed than their surroundings because the nonlinearity inevitably places some power there, but still suppressed relative to baseline. When these frequencies are not part of the target, the associated power does not contribute to performance and should be suppressed.

**Figure SI.2:**
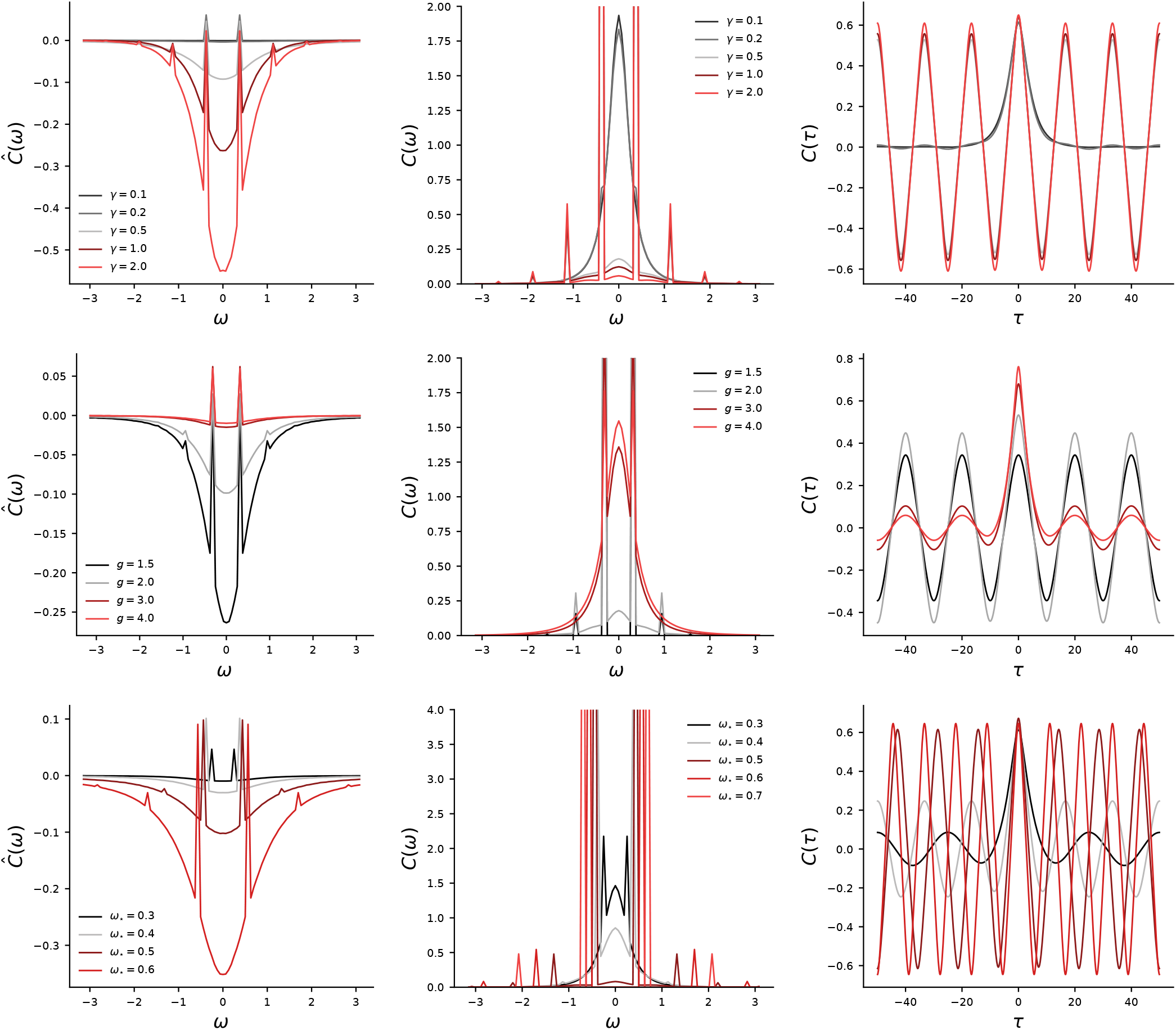
Time-translation-invariant DMFT solutions for autonomous (*C*^*I*^ = 0) nonlinear RNNs with *ϕ*(·) = tanh(·), trained to fit a Dirac target power spectrum *C*^*y*^(*ω*) = *δ*(*ω* − *ω*_⋆_). Columns show, from left to right, the conjugate order parameter *Ĉ*(*ω*), the power spectrum *C*(*ω*), and the time-domain autocorrelation *C*(*τ*). Each row varies one parameter while holding the other two fixed. Top row: varying *γ* with (*g, ω*_⋆_) = (2.5, 0.375). Middle row: varying *g* with (*γ, ω*_⋆_) = (0.5, 0.375). Bottom row: varying *ω*_⋆_ with (*γ, g*) = (0.5, 2.5). In the left column, *Ĉ*(*ω*) is positive near the target frequency *ω*_⋆_ (amplifying that frequency) and can become negative at other frequencies (suppressing task-irrelevant spectral content, including higher harmonics generated by the nonlinearity).

### SI.4 Comparing notions of effective dimensionality of activity

Over the course of this paper, we use two complementary notions of the effective dimensionality of activity. The starting point for both measures is the participation ratio of the finite-size autocorrelation *C*_*N*_ (*t, t*′) over a fixed observation window [−*T/*2, *T/*2],

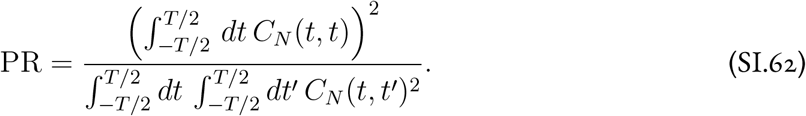

Here, we detail how the two measures we use arise from taking the limits *N* → ∞ and *T* → ∞ in different orders. What we will find is that if we first take *N* → ∞ and then *T* → ∞, the participation ratio is extensive in *T*, while if we first take *T* → ∞ and then *N* → ∞, the participation ratio is extensive in *N*.

***N* → ∞, then *T* → ∞**. If we take *N* → ∞ for fixed *T* and then take *T* → ∞, all fluctuations in the autocorrelation vanish, and we have *C*_*N*_ (*t, t*′) → *C*(*t, t*′) (almost-surely uniformly). Then, suppose that we take *T* ≫ 1, such that the system relaxes into a stationary state with *C*(*t, t*′) = *C*(*t* − *t*′). Assuming the contribution of the relaxation timescale to the integral is negligible, we then have

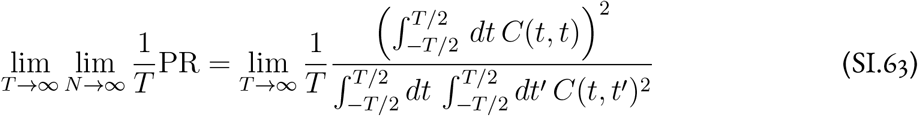

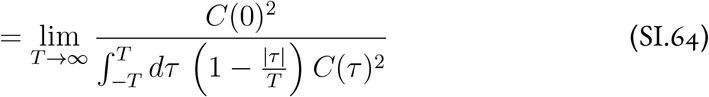

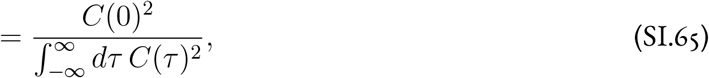

where we assume that *C*(*τ*) is bounded from above and is square-integrable, and moreover that |*τ* |*C*(*τ*)^2^ is integrable. Using the Fourier transform convention

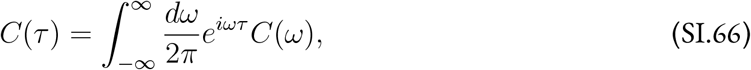

Parseval’s theorem implies that

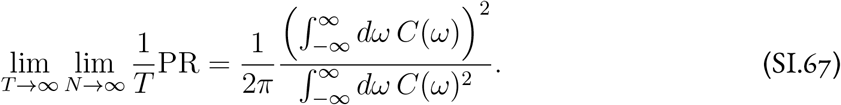

***T* → ∞, then *N* → ∞**. If *T* ≫ *N*, then we can no longer neglect finite-size fluctuations in *C*_*N*_. Assuming again that we are nearly in a stationary state, we decompose the finite-size correlation function as

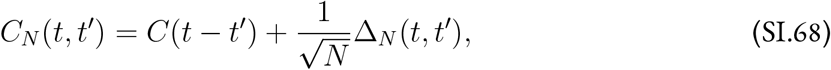

where Δ_*N*_ (*t, t*′) is an 𝒪_*N*_ (1) fluctuation term. Dividing both by *T* ^2^, the numerator and denominator of the participation ratio are then

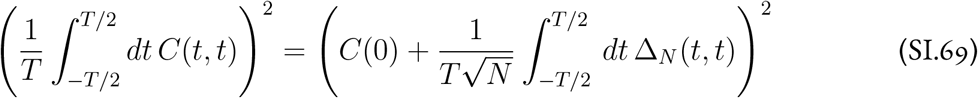

and

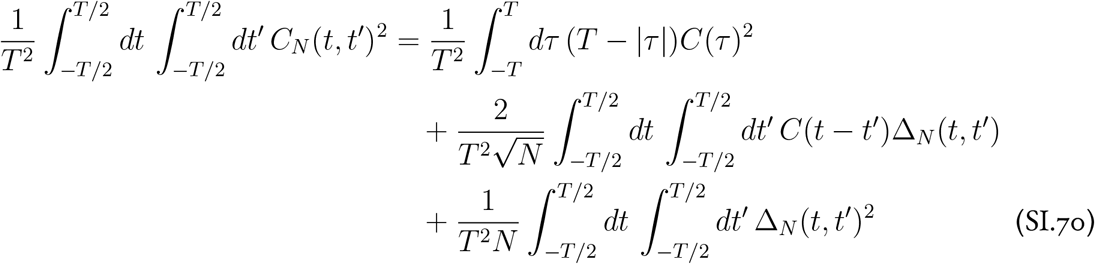

respectively. We now take *T* → ∞. The Δ_*N*_-dependent term in the numerator is negligible, as it has zero mean (over realizations) and standard deviation of order 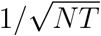. In the denominator, the contribution of the stationary covariance *C*(*τ*)^2^ is of order 1*/T* and is thus negligible. The second term in the denominator has mean zero and standard deviation of order 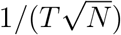 assuming that *C*(*τ*) is square-integrable, and is therefore also suppressed. Finally, the third term in the denominator is, up to negligible corrections, the order-one mean-square fluctuation amplitude

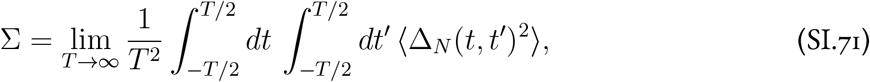

which is computable as in Clark et al. [26, 91]. This term dominates the denominator, leading to

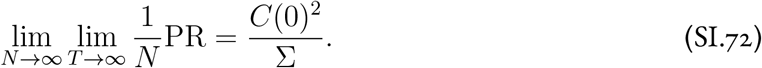

### SI.5 Sine wave (temporal generalization) task

We trained networks to autonomously generate the two-dimensional sinusoidal target

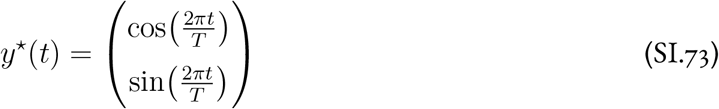

with period *T* = 10 (in units of *τ*), starting from a uniform initial condition *x*^0^ = 1 for all neurons, with no external input. We set *τ* = 1 throughout this task, so all times are dimensionless.

For each (*g, γ*) configuration, we trained 10 independent networks of size *N* = 2500 via Langevin gradient flow with inverse temperature *β* = 2000, step size 0.05, and 75,000 iterations; the Euler integration step was Δ*t* = 0.5. After training, networks were run autonomously for 5*T* to assess generalization and convergence to limit-cycle dynamics. We identified the onset of nonchaotic behavior by checking whether the normalized autocorrelation returned to unity (threshold 0.98) after 3*T* of autonomous evolution [54]. The DMFT equations were solved numerically on a denser (*g, γ*) grid using the alternating iteration procedure described in Sec. SI.1.3, with solutions warm-started across adjacent *γ* values to accelerate convergence. RNN simulations were run on a subset of 9 of the 20 *g* values; the DMFT was solved on the full grid. All parameters are given in Table SI.1.

### SI.6 Reaching task and neural data analysis

We used the dataset from Churchland et al. [6], as reanalyzed by Sussillo et al. [27], consisting of trial-averaged neural and EMG recordings from monkey J performing *B* = 27 reaching movements.

The dataset contains two distinct components. The first is the neural data, namely simultaneously recorded firing rates from 161 neurons in M1/PMd, which we use for the RNN–neural comparisons described below. The second is the RNN training data, consisting of precomputed inputs and target outputs from Sussillo et al. [27]. The inputs are seven-dimensional. The first six are the top principal components of preparatory neural activity, each with the same time-dependent modulation (ramping on during the preparatory period and turning off before movement onset). The seventh is a condition-independent hold cue that is on during the delay and then turns off to trigger the movement. The target outputs are 8-channel EMG signals that have been set to exactly zero until shortly before movement onset.

All data were originally provided at 10 ms resolution. We downsampled by a factor of 2 to a 20 ms time step, yielding 148 time bins for the RNN training data and 118 time bins for the neural recordings, with movement onset at bin 107 and bin 77 respectively. The provided inputs include 8 preparatory delay durations used by Sussillo et al. [27]; we used only zero delay. In the RNN, we set *τ* = 1 and Δ*t* = 0.4 in simulation units; because each integration step corresponds to one 20 ms data bin, the physical time constant is *τ*_phys_ = (20 ms)*/*(Δ*t/τ*) = 50 ms, consistent with the value used by Sussillo et al. [27]. Setting *τ* = 50 ms with Δ*t* = 20 ms as reported in the main text and Table SI.2 is equivalent, since only the ratio Δ*t/τ* matters.

EMG targets were shifted to have zero mean during the pre-movement baseline, and both inputs and EMG targets were rescaled to have approximately order-one magnitude when nonzero. To match the initialization conventions of Sussillo et al. [27], who included a factor of 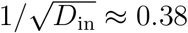 into the input weights at initialization, we further scaled the order-one inputs by a factor of 0.4.

We used *N* = 1024 neurons, compared to *N* = 300 in Sussillo et al. [27], to be in the large-*N* regime described by the DMFT.

Training was performed via Langevin gradient flow with gradient norm clipping (threshold 10, triggered extremely infrequently). Full parameters are given in Table SI.2.

#### Equilibration diagnostics

We verified that the Langevin sampler reached equilibrium through several diagnostics. Training MSE curves decrease and plateau before the end of sampling (Fig. 4B), with similar behavior for the full energy (including the Frobenius-norm terms). We also verified that individual weight matrix elements fluctuated on a timescale much shorter than the total sampling time (Fig. SI.3). We note that the MSE continued to decrease slightly when plotted on a log scale, particularly for small *g*. However, the eigenvalue spectrum of 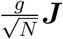, which is self-averaging in the large-*N* limit, showed negligible change over 100,000 iterations deep into sampling (Fig. SI.4), confirming that the macroscopic properties of the network had stabilized.

#### RNN–neural comparisons

We compared RNN population-level activity to simultaneously recorded M1/PMd firing rates in a ±400 ms window around movement onset (±20 time bins at 20 ms resolution). We considered two similarity metrics, centered kernel alignment (CKA) [56] and singular vector canonical correlation analysis (SVCCA) [92] with 10 components; and two preprocessing choices, z-scoring each unit or mean-centering only, yielding four comparisons in total. Sussillo et al. [27] used SVCCA with 10 components. For CKA, we computed the score between the *N* × (*B* × *n*_*T*_) RNN activity matrix and the corresponding 161 × (*B* × *n*_*T*_) neural activity matrix, where *n*_*T*_ = 40 is the number of time bins in the comparison window; SVCCA was computed analogously. Results are qualitatively similar across all four metric–preprocessing combinations (Fig. SI.5), with an intermediate degree of recurrent restructuring substantially improving the match to neural data relative to the reservoir limit. All four comparisons show clear nonmonotonicity in *γ* for most values of *g*, with the exception of CKA without z-scoring.

**Figure SI.3:**
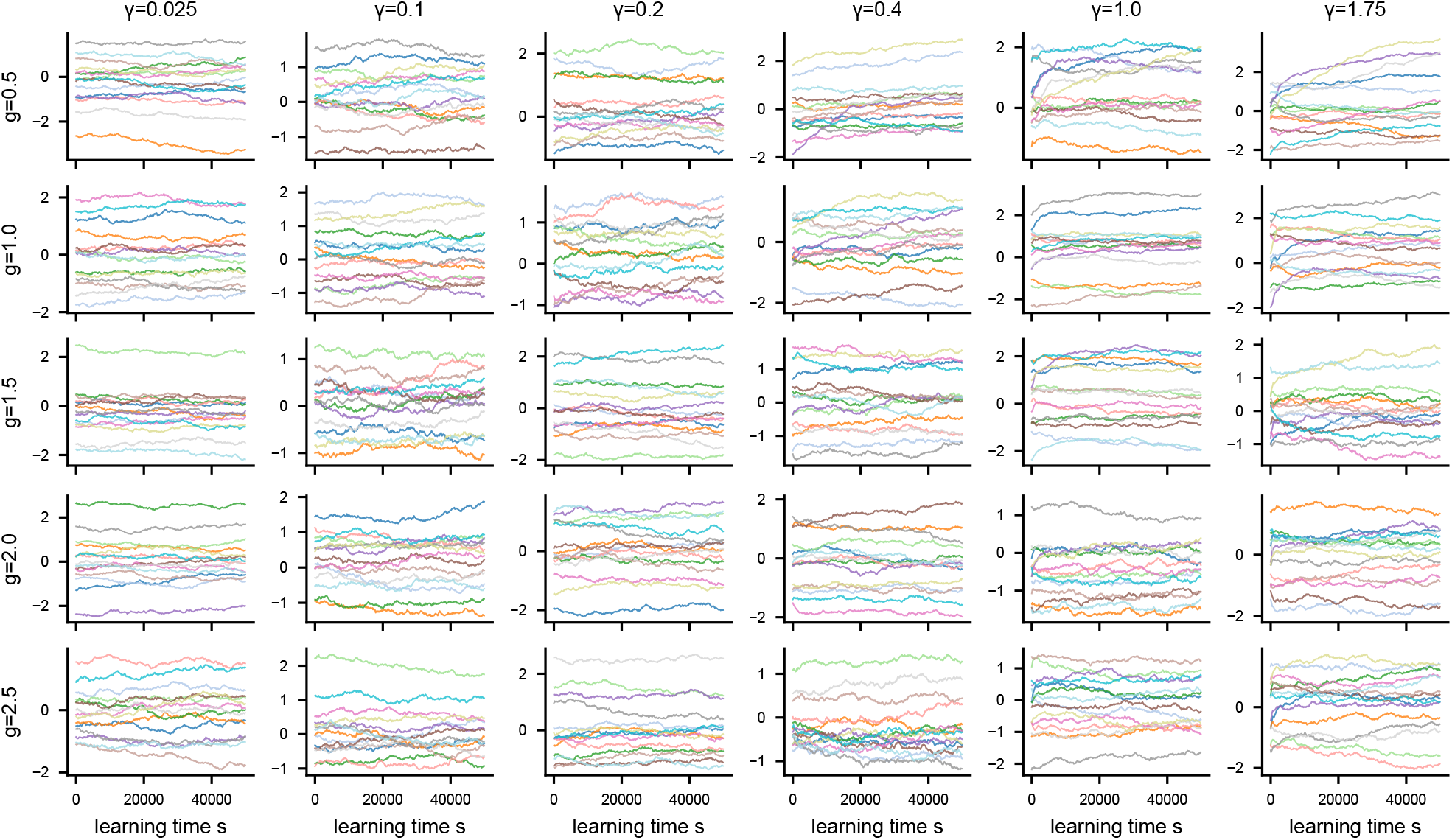
Weight traces during sampling. Traces of representative elements of *J*_*ij*_ over the course of Langevin gradient flow on the reaching task for a grid of *g* (rows) and *γ* (columns).

### SI.7 p-body generalization of the model

In addition to the RNN with two-body interactions between nodes, we can consider a *p*-neuron interaction generalization of the dynamics, reminiscent of *p*-spin models in the spin-glass literature [93]. These models have also gained some attention as descriptions of many-neuron interactions in biology, though the underlying physiology remains a subject of some debate [94]. The dynamics are

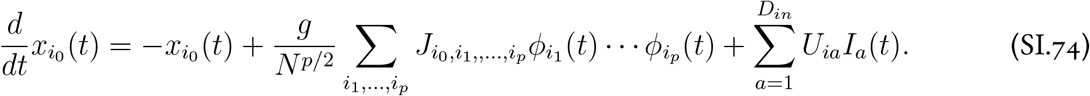

Under Langevin dynamics on the tensor ***J***, the read-in weights ***U***, and the readout weights ***V***, we arrive at a structurally identical DMFT in terms of the autocorrelation 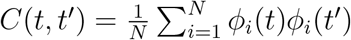, In the *β* → ∞ limit, the DMFT action takes the form

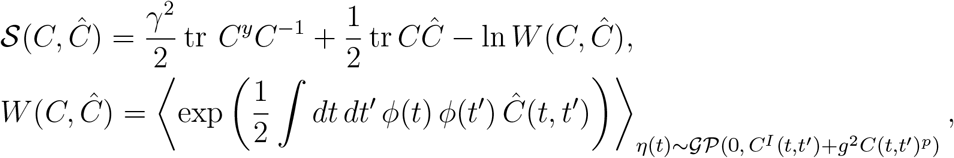

**Figure SI.4:**
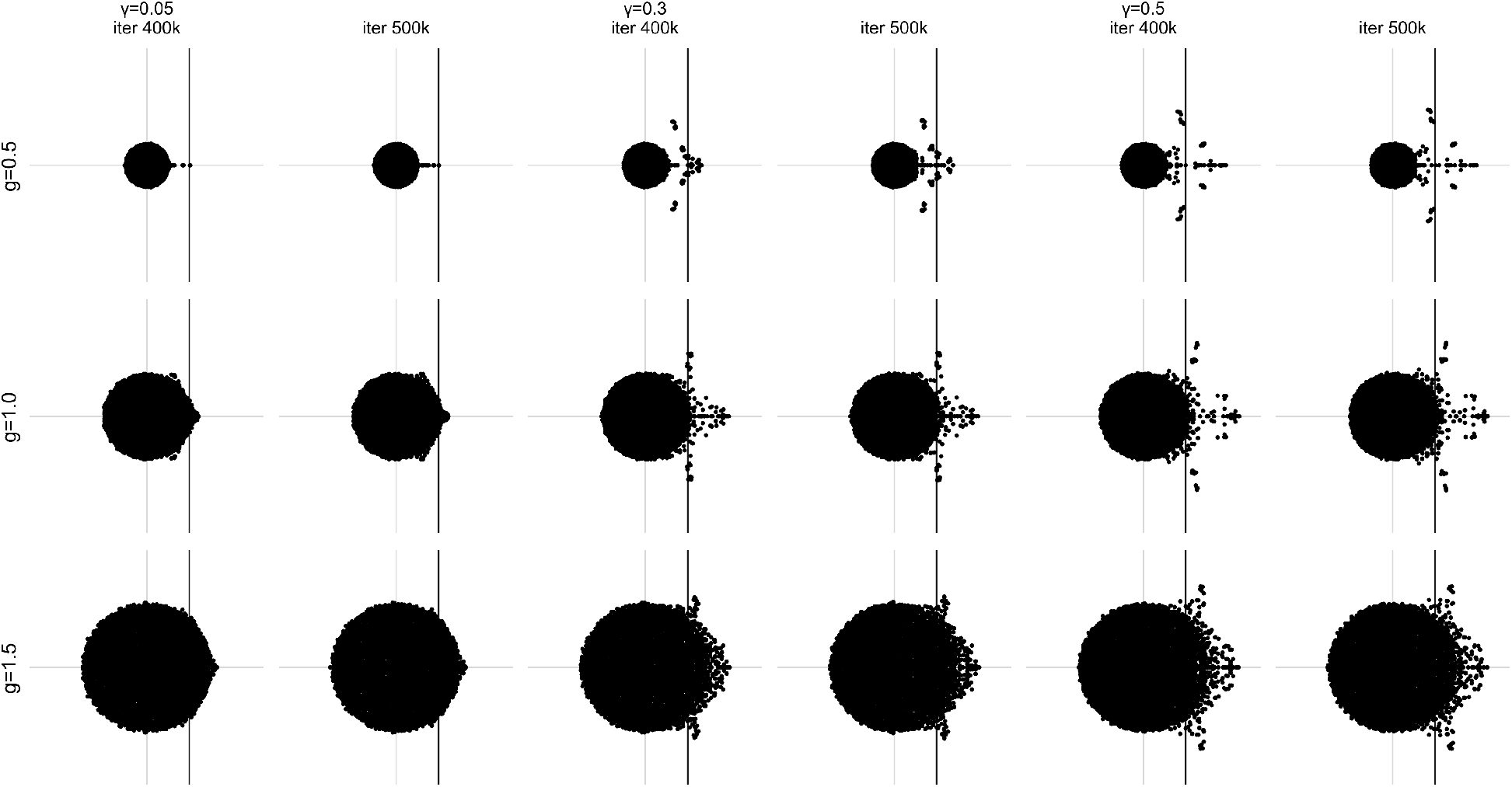
Eigenvalue spectra are stable across sampling. Eigenvalue spectra of 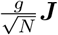 at iteration 400,000 (left in each pair) and iteration 500,000 (right), for selected (*g, γ*) configurations. Rows correspond to *g*; column pairs correspond to *γ*. Spectra show negligible change between the two checkpoints.

The main modification relative to the *p* = 1 case is that the noise driving the internal recurrent dynamics *η*(*t*) has covariance *g*^2^*C*(*t, t*′)^*p*^ instead of *g*^2^*C*(*t, t*′). As before, the *N* → ∞ limit is determined by the saddle-point equations 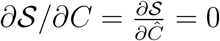.

#### Nonlinear Dynamics with Linear Activations

One advantage of this model is that the dynamics can still be chaotic and autonomous even with linear activations *ϕ*(*x*) = *x*, provided that *p >* 1. For this choice, the single-site distribution for *x* remarkably remains Gaussian, even in the feature-learning regime. The DMFT equations for *C* and *Ĉ* therefore close without any single-site average (shown here in the *β* → ∞ limit),

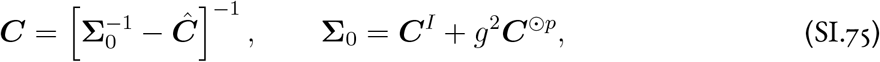

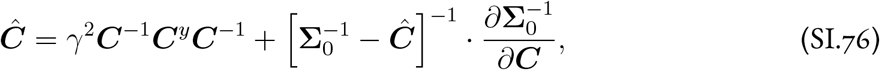

where ***C***^⊙*p*^ denotes the elementwise *p*th power. This model’s DMFT could therefore in principle be solved without resorting to Monte Carlo integration. To maintain stability in the chaotic regime without external drive (*C*^*I*^ = 0) and without a saturating nonlinearity, one can constrain the dynamics of the norm of *x* (restricting to the sphere) [93]. We leave detailed analysis of this model in the feature-learning regime to future work.

**Figure SI.5:**
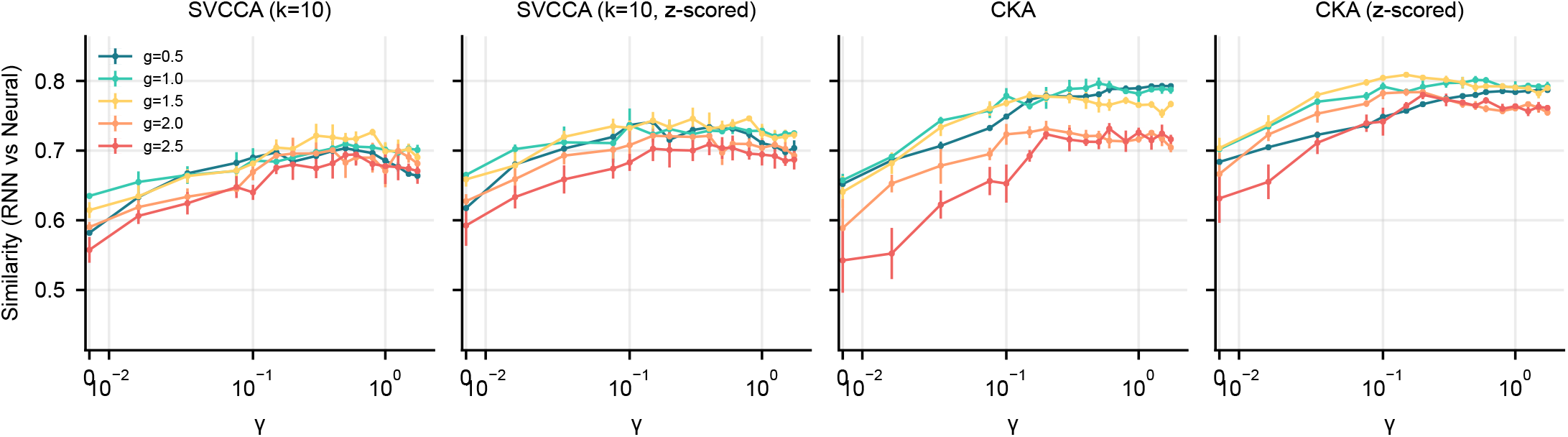
RNN–neural similarity across metrics and *z*-scoring choices. Similarity between RNN and M1/PMd population-level activity as a function of *γ* for different values of *g* (colors), computed in a ± 400 ms window around movement onset. The rightmost panel is the same as Fig. 5A.

**Table SI.1:** Sine wave task parameters. RNN architecture and Langevin gradient flow training (top), hyperparameter grid (middle), and DMFT numerical solver settings (bottom).

| Parameter | Symbol | Value |
| --- | --- | --- |
| *Architecture* |  |  |
| Neurons | $N$ | 2500 |
| Activation function | $\phi(\cdot)$ | $\tanh(\cdot)$ |
| Time constant | $\tau$ | 1 (dimensionless) |
| Euler step | $\Delta t$ | 0.5 |
| Output dimensions | $D_{\text{out}}$ | 2 (cos, sin) |
| External inputs |  | None |
| Initial condition | $x^0$ | 1.0 |
| Target period | $T$ | 10 |
| *Langevin gradient flow training* |  |  |
| Inverse temperature | $\beta$ | 2000 |
| Step size |  | 0.05 |
| Training iterations |  | 75,000 |
| Gradient clipping |  | None |
| Weight initialization | $J_{ij}, V_{ia}$ | $\mathcal{N}(0, 1)$ i.i.d. |
| Independent runs | | 10 per $(g, \gamma)$ |
| *Hyperparameter grid* |  |  |
| Recurrent gain | $g$ | 20 values in $[1.0, 3.0]$ |
| Output coupling | $\gamma$ | 27 values in $[0, 1.2]$ ($g \leq 1.5$);<br>19 values in $[0, 1.2]$ ($g > 1.5$) |
| Trained subset | | 9 of 20 $g$ values |
| *DMFT solver* |  |  |
| Outer iterations |  | 300 |
| Inner $C$-update steps | | 100 |
| MC samples ($C$ update) | | $10^5$ |
| MC samples ($\hat{C}$ update, action gradient) | | $10^6$ |
| Cholesky regularization | | $10^{-5}$ |
| $C$ mixing rate | | 0.5 |
| $\hat{C}$ mixing rate | | 0.1 |
| $C$ initialization | | Identity |
| $\hat{C}$ initialization | | Zeros |
| Warm-start across $\gamma$ | | Yes |

**Table SI.2:** Motor cortex RNN parameters. Architecture, training, and hyperparameter grid for the reaching task of Sec. 2.6. Each (*g, γ*) pair was trained with 5 independent random initializations, for a total of 375 networks.

| Parameter | Symbol | Value |
| --- | --- | --- |
| *Architecture* |  |  |
| Neurons | $N$ | 1024 |
| Activation function | $\phi(\cdot)$ | $\tanh(\cdot)$ |
| Time constant | $\tau$ | 50 ms |
| Euler step | $\Delta t$ | 20 ms |
| Input dimensions | $D_{\text{in}}$ | 7 (6 temporally-modulated preparatory PCs + 1 go cue) |
| Output dimensions | $D_{\text{out}}$ | 8 (EMG activity) |
| Initial condition | $x^0$ | 0.0 |
| Training conditions | $B$ | 27 reach directions |
| *Langevin gradient flow training* |  |  |
| Inverse temperature | $\beta$ | $10^6$ |
| Step size |  | 0.1 |
| Training iterations |  | 500,000 |
| Gradient clip norm |  | 10 |
| Weight initialization | $J_{ij}, U_{ia}, V_{ia}$ | $\mathcal{N}(0, 1)$ i.i.d. |
| *Hyperparameter grid* |  |  |
| Recurrent gain | $g$ | {0.5, 1.0, 1.5, 2.0, 2.5} |
| Output coupling | $\gamma$ | 15 values in [0.025, 1.75] |
| Independent runs | | 5 per $(g, \gamma)$ |
| Total networks |  | 375 |

## Footnotes

1 Code and data are available at https://github.com/davidclark1/RNN-Learning-Theory.

2 A subset of the authors presented preliminary results at the 2024 Computational and Systems Neuroscience Conference [37], including the general dynamical mean-field formulation and the analytical solution for linear RNNs.

